# Advancing the global statistical standard for urban ecosystem accounts

**DOI:** 10.1101/2025.03.05.641195

**Authors:** Javier Babí Almenar, Chiara Cortinovis, Sara Vallecillo, Davide Geneletti, Balint Czucz, Federica Marando, Grazia Zulian, Anna M Addamo, Alessandra La Notte, Renato Casagrandi

## Abstract

The System of Environmental-Economic Accounting-Ecosystem Accounting (SEEA-EA), adopted by UNSD, provides a standardized global framework for measuring and monitoring ecosystems’ extent, condition, and services. However, its application to urban ecosystems faces conceptual and operational challenges. Building on SEEA-EA, we propose advancing the framework for thematic urban ecosystem accounting, identifying main challenges and framing potential solutions based on existing lessons and approaches. Through a literature review on ecosystem accounting and urban science, we identified 24 challenges, with lessons and approaches suggested for 17 of them. Results show that many challenges are highly interconnected and shared with accounts for other ecosystem types. Urban-specific challenges include a lack of consensus in defining urban ecosystems, their specific assets, and their classifications. Additionally, findings highlight the need for defining appropriate methods to capture socio-ecological degradation, impacts, and dependencies of urban ecosystems. Suggested solutions include adapting the accounting structure and prioritizing the resolution of urban-specific challenges.

## 1. Main

Biodiversity and healthy ecosystems are cornerstones of human health and well-being, underpinning ecosystem service supply. Yet, both continue to decline^1–5^. It is therefore crucial to track changes in the status of ecosystems and biodiversity, along with their societal and economic impacts, in a regular, consistent, and comparable manner. Ecosystem accounts provide a structured framework for measuring and monitoring the extent and condition of ecosystems, their service flows, and their value as assets^6,7^. Their implementation is expected to support policymaking across sectors and governance levels, from international to local^8–10^. Recognizing the value of ecosystem accounts to inform sustainable ecosystem management, countries are increasingly adopting them through legislation^11,12^.

Ecosystem accounts may be applied to any ecosystem type, including anthropogenic ones. Urban ecosystems - characterized by the presence of people and a mix of natural and artificial features^13^ - are particularly relevant from an ecosystem accounting perspective. As primary habitats for humans and hotspots of ecosystem services’ demand, urban ecosystems are key drivers of global and local biodiversity changes^14–18^. A global framework for urban ecosystem accounts would be a valuable instrument for the still-emergent urban science-policy-practice networks^19^, helping to translate knowledge into informed actions that address biodiversity loss and ecosystem degradation.

The System of Environmental-Economic Accounting Ecosystem-Accounting (SEEA-EA), adopted in 2021 by the UN Statistical Commission, is the globally recognized statistical standard for ecosystem accounts^13^. SEEA-EA is a spatially-based integrated statistical framework applicable to all ecosystem types, including urban ecosystems. It comprises five core accounting tables: 1) ecosystem extent, 2) ecosystem condition, 3) ecosystem service flows in biophysical terms, 4) ecosystem service flows in monetary terms, and 5) monetary ecosystem assets. The first three are adopted as part of the statistical standard, while the developments for the last two are currently recognized only as statistical principles and recommendations^13,20^. A glossary of key SEEA-EA terms is provided in the Supplementary Information (SI)1.

Although SEEA-EA may be applied to compile urban ecosystem accounts, the current scientific conceptualisation of urban ecosystems conflicts with some of SEEA-EA’s core principles. For instance, SEEA-EA requires ecosystem assets to represent ecosystems and be mutually exclusive conceptually and geographically^13^. However, as conceptualised by urban and landscape ecologists, urban ecosystems are heterogeneous landscape mosaics of fine-grained patches (ecosystem assets) representing multiple ecosystem types, with boundaries that are often fuzzy^21,22^. Additionally, ecosystem condition in urban ecosystems is shaped not only by natural abiotic and biotic components but also by social and technological factors^23,24^, which SEEA-EA does not currently account for^25^. Consequently, degradation in urban ecosystems is not solely ecological but also socio-ecological. Failing to acknowledge these aspects when developing a global framework for urban ecosystem accounting risks creating a framework perceived as inadequate or illegitimate by urban ecologists and related science-policy-practice networks, ultimately leading to a misuse.

To overcome such inconsistencies, SEEA-EA encourages the development of thematic accounts for urban ecosystems^13^. Thematic accounts provide greater flexibility than traditional SEEA-EA accounts, hereafter, referred to as SEEA-EA general accounts. However, the UN Statistical Commission has yet to formally adopt thematic accounts as part of the statistical standard or recognize them as part of formal principles and recommendations^13,20^. In fact, thematic urban ecosystem accounts remain underdeveloped as a standard, potentially leading to inconsistencies with SEEA-EA general accounts and the proliferation of multiple incompatible local or regional frameworks^26^. This issue threatens the ability of SEEA-EA to support sustainable urban ecosystem management in a globally coherent way.

Thus, developing a global framework for urban ecosystem accounting in the form of thematic SEEA-EA accounts is essential. This framework must integrate seamlessly with SEEA-EA general accounts and enable a unified, systematic approach for urban ecosystem accounting. Achieving this requires identifying and addressing remaining challenges posed by SEEA-EA when applied to urban ecosystems.

This paper aims to advance the development of the global framework for urban ecosystem accounting, further standardizing SEEA-EA thematic accounts by:

i. Identifying key conceptual and operational challenges for thematic urban ecosystem accounts. Conceptual challenges pertain to unresolved theoretical issues, while operational challenges involve practical gaps or issues that should be addressed to establish clear procedures.
ii. Presenting lessons learnt and approaches from past experiences to inform potential solutions. Lessons point either to univocal solutions or to key constraints when addressing specific challenges, while approaches represent potential alternatives that occasionally appear contradictory in the literature.

Challenges, approaches and lessons were identified through a review of key SEEA-EA documents, followed by a systematic review of both grey and scientific literature on ecosystem accounting. These reviews were supplemented by a thematic review of scientific literature on challenges relevant beyond accounting.

## 2. Results

The review encompassed 130 documents, including 87 from the systematic review on ecosystem accounting, of which 50 were case studies. These case studies were conducted within 19 countries at local, regional, or national levels, with some in Europe developed at a transnational level. Only studies from six countries specifically addressed urban ecosystem accounts (Fig. 1). Across these documents, 24 challenges were identified (Table 1), 13 of which are conceptual. Lessons and approaches were identified for 18 challenges (Table 2).

**Fig. 1.**
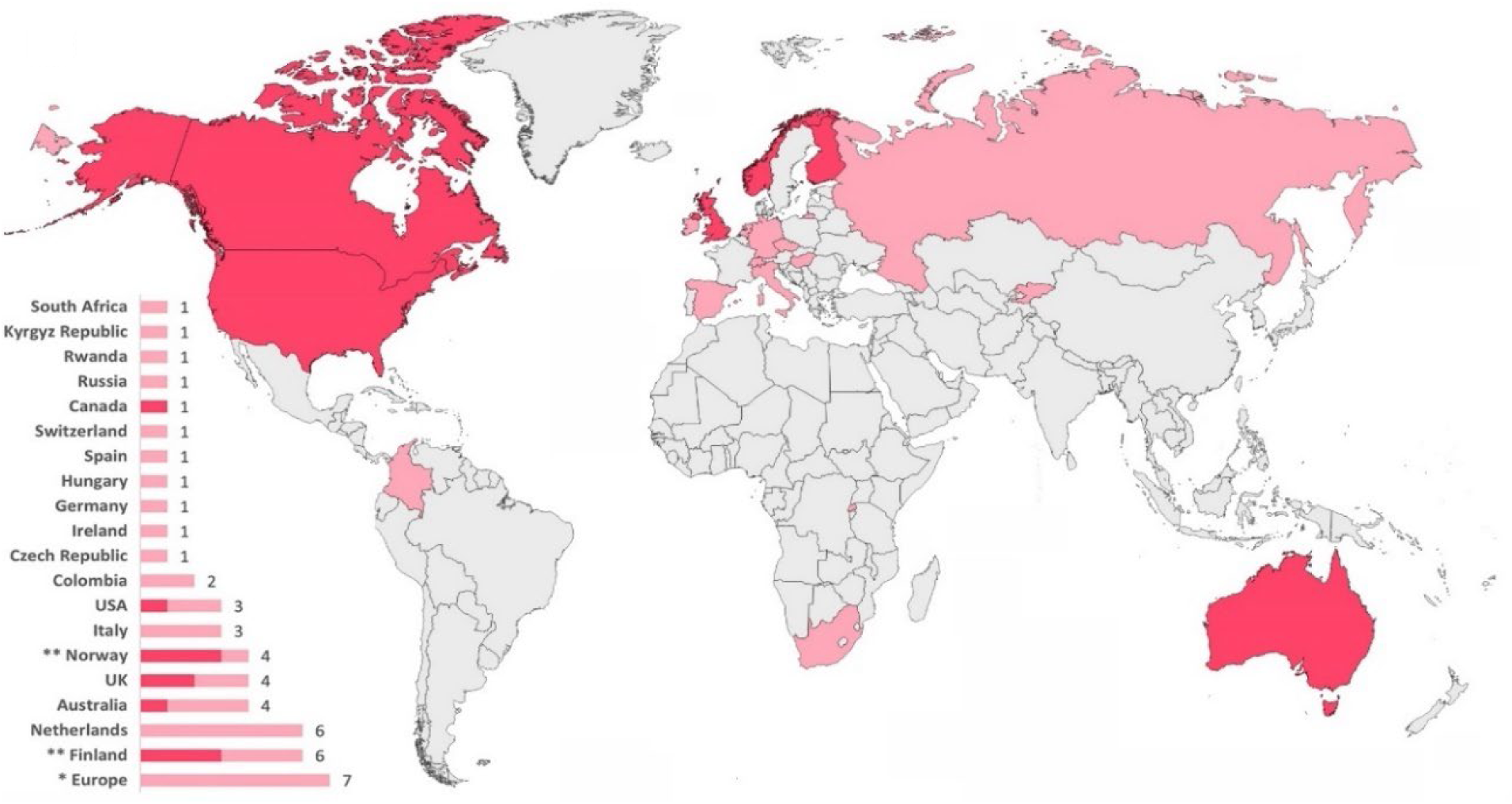
Case studies of ecosystem accounts (light red), differentiating those focused exclusively on urban ecosystem accounts (red). World map showing countries where ecosystem accounts were tested in the reviewed literature. The bar chart displays the number of case studies per country. The case studies applied SEEA-EA, its previous experimental version, or, in a few cases, very similar ad-hoc frameworks that referred to SEEA-EA. **Note:** *Case studies of ecosystem accounting developed at European level, including 25 or more countries. **One document includes three case studies from Finland and one from Norway, which are counted as independent cases.

**Table 1.** Challenges of thematic urban ecosystem accounting, grouped by SEEA-EA account table, relevant ecosystem types (classified as U: urban; AE: anthropogenic ecosystems, including urban ecosystems; MAE: Most or All Ecosystems), and challenge types (C: Conceptual; O: Operational). Additional details for each challenge are provided in Supplementary Information (SI3).

| Account Table | Ecosystem Types | Challenge Type | Id | Short description | Refs. |
| --- | --- | --- | --- | --- | --- |
| Extent | U | C | 1 | There is still no univocal distinction between urban ecosystem assets and artificial surfaces. | 27–34 |
|  | U | C | 2 | There is still no univocal procedure for the spatial delineation of urban ecosystems and their accounting areas. | 13,26–28,35–44 |
|  | U | C | 3 | At a global level, urban ecosystems still lack an ecologically sound, agreed-upon classification of homogeneous ecosystem areas, along with the principles to develop it. | 30,42–47 |
|  | U | C | 4 | Urban ecosystems still lack a common ecologically sound classification of the fine-scale patches (assets) that compose them. | 28,30,42,44,46,48–52 |
| Condition | AE | C | 5 | Ecosystem condition variables offer just a partial representation of socio-ecological-technological characteristics driving the functioning and quality of urban ecosystems. | 23,24,53 |
|  | AE | C | 6 | There is no clear conceptual basis for defining reference conditions in anthropogenic ecosystems, being “ecological integrity” inadequate as a default concept for them. | 54–56 |
|  | AE | C | 7 | Ecosystem condition accounts cannot fully capture socio-ecological degradation and indirect degradation in urban ecosystems. | 44,57–61 |
|  | MAE | O | 8 | There are difficulties establishing boundaries between ecosystem condition and ecosystem extent accounts, particularly at fine levels of ecosystem classification. | 38,62 |
|  | MAE | O | 9 | There is no consensus yet on a common minimum set of ecosystem condition variables and on their reference levels. | 13,24,42,54,63–65 |
|  | MAE | O | 10 | There is still insufficient evidence and consensus on the appropriate procedure(s) to aggregate ecosystem condition indicators into indices. | 64 |
| Services | MAE | C | 11 | There is no consensus about the concept of ecosystem capacity, and its connection to extent and condition and to other related concepts (e.g. degradation, sustainability, and resilience). | 13,63,66–71 |
|  | MAE | C | 12 | Ecosystem service accounts fall short in informing whether there are mismatches in service flows that represent sustainability issues (e.g. degradation), unmet societal needs, and/or missed flows. | 58,61,65–68,72–75 |
|  | MAE | C | 13 | There is still missing a complementary pluralistic valuation of ecosystem services (and assets) that represents multiple systems of values. | 13,24,53,59,76–82 |
|  | MAE | O | 14 | There is the need for an explicit consideration of intermediate ecosystem services, along with clear rules for allocating inter- and intra-ecosystem service flows and for avoiding double counting. | 65,75,83,84 |
| Monetary Assets | MAE | C | 15 | Modelling future ecosystem service flows entails interlinked, difficult-to-make assumptions, and inherits the uncertainty of the data from other account tables, which is not reflected in monetary ecosystem asset accounts and their methodological framework yet. | 13,26,56,70,76,85–87 |
|  | MAE | C | 16 | Robust agreed-upon principles for estimating and allocating value loss due to ecosystem degradation are still missing in monetary ecosystem asset accounts and its methodological framework. | 56,58,61,70,74,76,85,88 |
|  | MAE | O | 17 | There is still not enough evidence and consensus on the appropriate “asset life” value(s) to be used in monetary ecosystem asset accounts. | 27,76,86,87 |
|  | MAE | O | 18 | There is still not enough evidence and consensus on the most appropriate discount rate(s) to be applied to future ES flows. | 27,70,76,86,87 |
| Cross-cutting | U | C | 19 | Varied and unclear policy uses of urban ecosystem accounts across multiple spatial levels (national, regional, local) create diverse requirements, risking loss of coherent framing. | 26,38–40,56,62,89–95 |
|  | MAE | O | 20 | There are still missing complete and consistent examples of ecosystem accounts that can serve as practical guidance. | 13,77,96,97 |
|  | MAE | O | 21 | Ecosystem accounts still lack an explicit assessment and reporting of uncertainties, along with its agreed-upon procedure. | 27,33,40,52,62,70,75,76,86,98–101 |
|  | MAE | O | 22 | Expertise, time, and resource demand generate practical barriers for the implementation of ecosystem accounts. | 59,62,69,77,97 |
|  | MAE | O | 23 | There are still gaps, and lack of consensus, regarding suitable input data, data quality frameworks, and generalizable models for use in ecosystem accounts. | 13,32,33,39,40,62,75,77,84,88,90,91,94,97,99,100,102–115 |
|  | MAE | O | 24 | There is still a need to establish common principles and practices for developing data infrastructures for ecosystem accounting, ensuring interoperability and data sharing among stakeholders. | 13,59,84,90,101 |

**Table 2.**
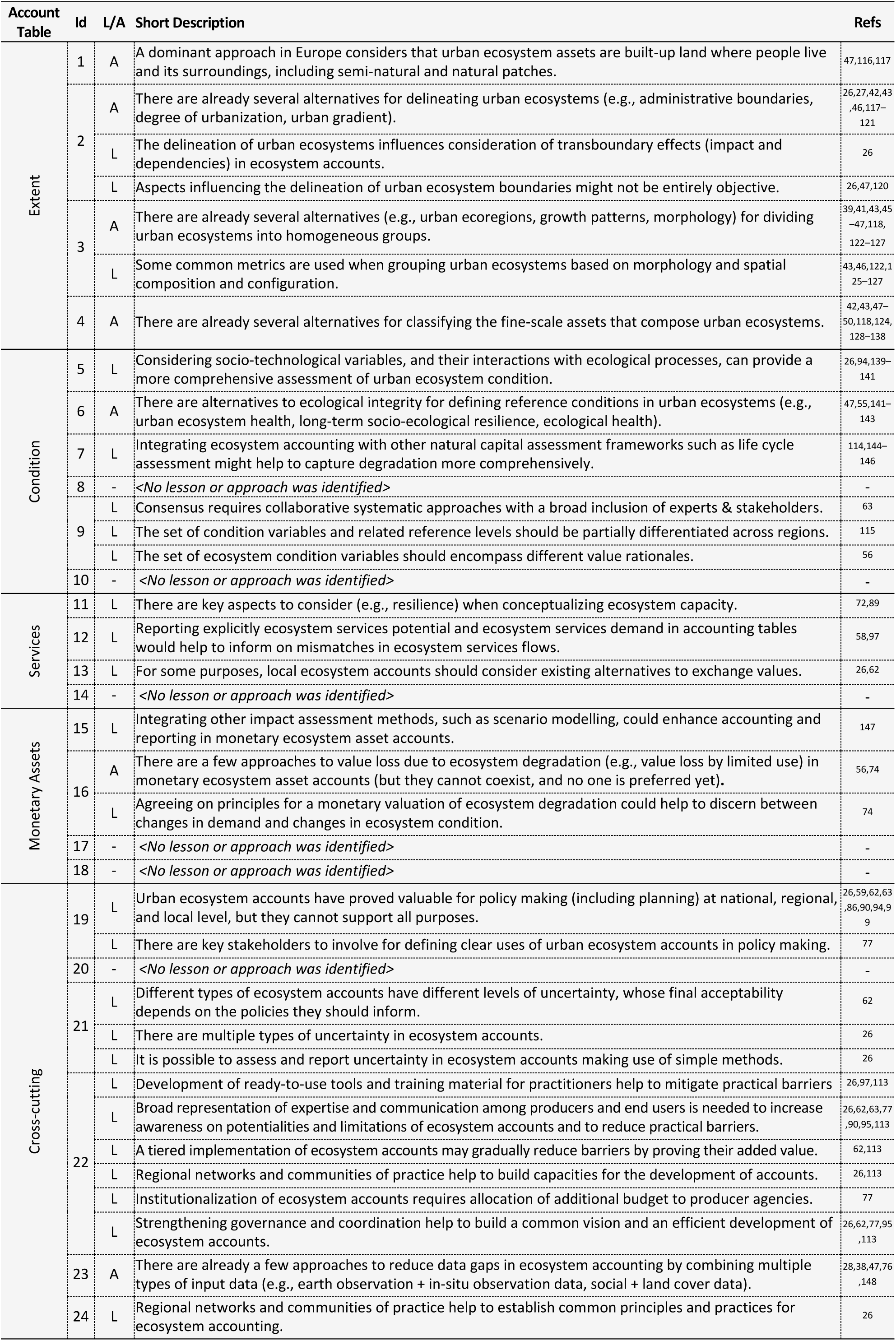
Lessons (L) and approaches (A) for thematic urban ecosystem accounting organized by account table and challenge type (challenge Ids as in. Table 1). Additional details for each L and A are provided in Supplementary Information (SI4).

Challenges specific to urban and anthropogenic ecosystems, the focus of the following sections, were identified exclusively within extent, condition, and cross-cutting categories. The latter refers to challenges relevant to multiple accounting tables. Further details on the data gathered, as well as descriptions of all identified challenges, approaches and lessons, are provided in SI.2–4.

### 2.1. Characterisation and classification of urban ecosystems and their fine-scale patches

The literature reviewed highlights a lack of consensus on the definition and characterization of urban ecosystems^27,28,31–34^. Specifically, the distinction between artificial surfaces and urban ecosystem assets remains ambiguous (Table 1 - Challenge [Ch.]1), with accounting exercises not always classifying the same set of features as urban ecosystem assets^28,31^. This variability partly arises from how the boundaries of urban ecosystem accounts are defined (Ch.2). While several approaches have been proposed to determine which artificial surfaces qualify as urban ecosystem assets, no consensus has been reached^47,116,117^. A key suggested criterion is that, to qualify as an urban ecosystem asset, artificial surfaces must be related to built-up areas where humans reside^47,116^. This criterion seeks to exclude isolated infrastructures (e.g., mining sites, energy-related facilities) that are not physically or functionally tied to specific human settlements.

There is also no agreed-upon ecologically sound classification system to divide urban ecosystems into groups of homogeneous ecosystem areas (Ch.3)^43,45–47^. Globally, urban ecosystems are often treated as a single homogeneous group^149^. However, they span multiple regions, biomes and climates, exhibiting different urban forms shaped by surrounding environments, historical legacies, and variations in socio-demographic, ecological, and technological characteristics (Fig. 2). Scholars are exploring potential classification approaches, such as those based on urban ecoregions, growth patterns, and morphology, to categorize urban ecosystems into multiple coherent groups of homogeneous ecosystem areas^39,43,45–47,118,122–127^.

**Fig. 2.**
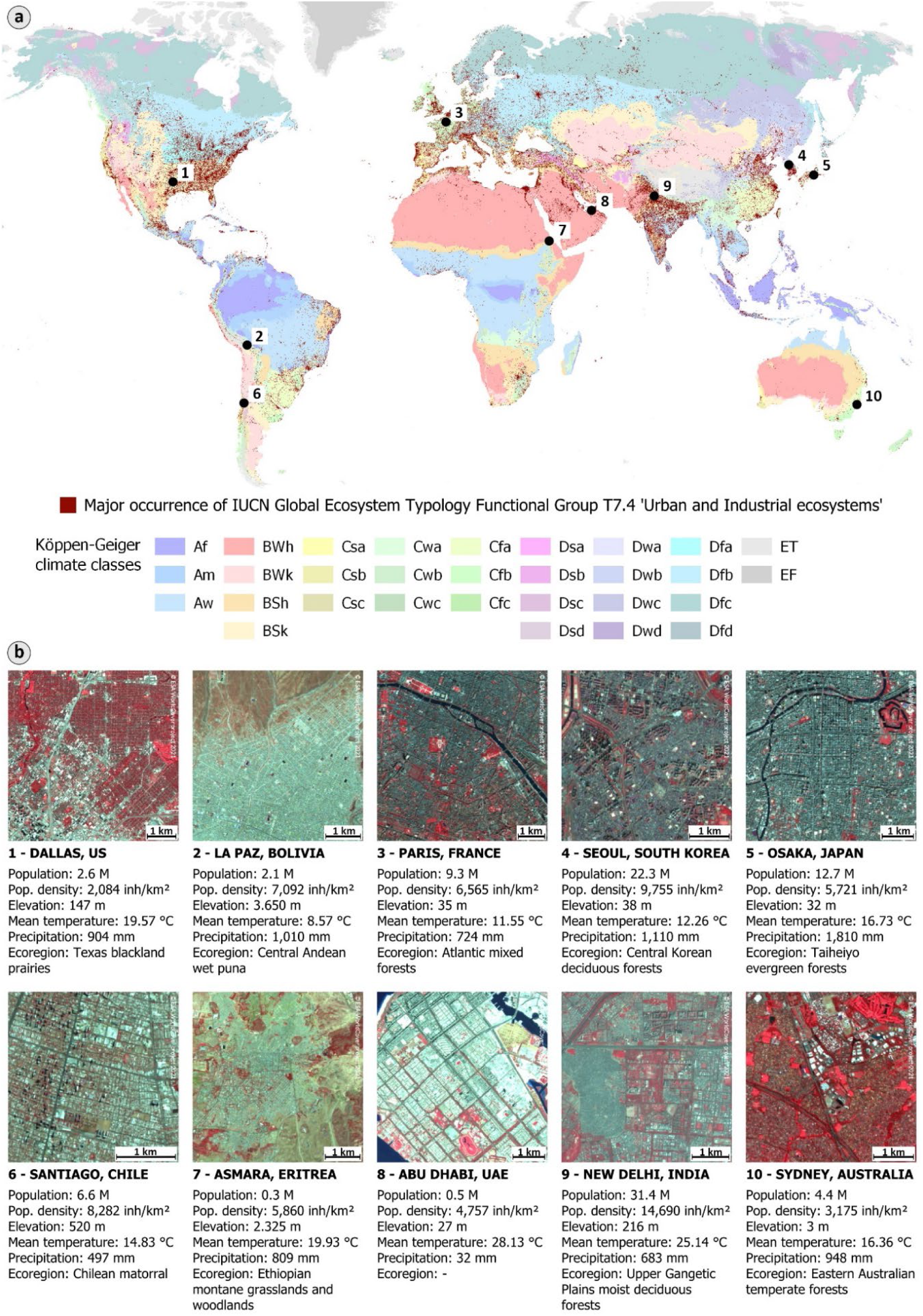
Visualization of the diverse climates, (eco)regions, and environmental and socio-demographic characteristics across which urban ecosystems extend worldwide, underscoring the relevance of classifying these systems into multiple groups of homogeneous ecosystem areas. **a)** Global distribution of urban and industrial ecosystems, as defined in the IUCN Global Ecosystem Typology149,153, overlaid on a global map of Köppen-Geiger climate classes154, illustrating the range of climates on which urban ecosystems are present. **b)** Aerial imagery snapshots (Sentinel-2 False Color Composite, 2021) of 10 urban ecosystems worldwide. In these illustrative snapshots, vegetation appears in red, while annotations highlight key factors such as population, environmental conditions, and ecoregion—underscoring the global diversity of urban ecosystems. The data for these factors come from the GHS Urban Centre Database R2024 of the Joint Research Centre, which delineates individual urban ecosystems based on the Degree of Urbanisation approach.

Similarly, there is no consensus on an ecologically sound classification for the fine-scale patches (assets) that compose urban ecosystems (Ch.4)^28,30,50,52,133,150^. Traditionally, these assets are classified based on land cover and land use^47,134,135^, but alternatives, such as local climate zones^28^, have been applied in some urban ecosystem accounting studies. Several scholars advocate for asset classifications (e.g., HERCULES^135^, STURLA^151^) that (i) avoid mixing land cover and land use, (ii) incorporate the third dimension (height), and (iii) treat interconnected artificial, natural, and semi-natural features as unified patches, preventing classification artifacts^134,135,151,152^. They argue that traditional classifications hinder a robust ecological understanding of how the composition and configuration of urban ecosystem assets influence ecological functions^135^, as well as condition, service supply and demand, and asset values.

Both classification issues (Ch.3,4), stem from a lack of scientific consensus, as evidenced by ongoing debates within urban ecology and related fields^42,50,51,134,135^. This explains why only approaches and lessons highlighting key considerations were identified. The absence of scientific consensus similarly affects challenges that are not urban-specific, such as estimating and allocating value loss due to ecosystem degradation (Ch.16), where fundamental issues remain debated within environmental and ecological economics^58,74,76^.

### 2.2. Tracking socio-ecological degradation, impacts and dependencies

Several limitations in condition accounts, combined with shortcomings in services and monetary ecosystem asset accounts, hinder the comprehensive tracking of urban socio-ecological impacts and dependencies.

For condition accounts, scholars highlight limitations in capturing: i) local degradation that is not strictly ecological; and ii) how urban ecosystems influence the degradation of nearby and far-away assets on which they depend, which may indirectly lead to urban degradation (Ch.7).

Regarding local degradation, limitations arise from excluding social and technological characteristics, leading to a partial representation of urban ecosystems (Ch.5). This omission may overlook certain types of socio-ecological degradation, such as declining public health or increasing inequalities in the distribution of ecosystem assets and services. Additionally, shortcomings may partly stem from reliance on the concept of ecological integrity to define reference conditions in urban ecosystems (Ch.6). Karr and colleagues^55^, the first a proponent of the concept of ecological integrity as referenced in SEEA-EA, note that ecological integrity, which exclude humans by definition, may not provide appropriate reference conditions (endpoints) for ecosystems under intensive human use, such as agroecosystems or urban ecosystems. Instead, they advocate for using ecological health as a more suitable concept^55^. Similar alternatives, such as urban ecosystem health, have been proposed by other authors^141,155,156^. These alternatives acknowledge humans as intrinsic components of the system, allowing characteristics of human populations and built infrastructure to be considered as condition variables^55,141,155^.

Some approaches also suggest that to capture how urban ecosystems drive the degradation of nearby or far-away ecosystem assets, assessments against a reference condition should account not only for local socio-ecological degradation but also for how urban characteristics influence service demand both locally and remotely^47,141,155^. These suggestions extend beyond standard SEEA-EA rules, which typically define ecosystem condition based on the overall ecological quality of a given ecosystem in terms of its local biotic and abiotic characteristics. In effect, these approaches highlight the strong telecoupling relationships between urban ecosystems and other ecosystems^114,144^, driven by social, economic, and technological factors^116,144^. They emphasize the importance of incorporating these connections when assessing urban ecosystem condition and its changes, particularly if condition accounts are intended to inform urban sustainability analyses. Some scholars suggested to use complementary natural capital assessment frameworks, such as life cycle assessment, to characterize telecoupling relations by tracking supply-chain connections^114,145,146^.

To better track socio-ecological degradation, impacts and dependencies of urban ecosystems, the literature also stresses the need for agreed-upon principles to estimate and allocate value loss from ecosystem degradation in monetary ecosystem asset accounts (Ch.16). In past studies, both ecosystem degradation and the explicit consideration of assumptions and uncertainties in modelling future ecosystem service flows (Ch.15) have often been overlooked when quantifying changes in asset values^86,87^. To minimise these issues, one paper suggests the use of scenario modelling^147^.

### 2.3. The relevance of policy uses for a coherent framework

The ability to track socio-ecological impacts and dependencies through thematic urban ecosystem accounts is also influenced by how urban ecosystems are delineated (Ch.2). Different approaches, such as using municipal boundaries or buffers around built-up areas, affect the inclusion or exclusion of specific ecosystem assets, thereby influencing the consideration of transboundary effects (i.e., impacts and dependencies)^26^. Some scholars note that delineating these boundaries is not entirely objective^26,120^, as it partly depends on the policy uses of the accounts. Indeed, the lack of clear policy uses is highlighted as a cross-cutting challenge (Ch.19), alongside a weak policy pull, an issue also faced by SEEA-EA general accounts^90^. For thematic urban ecosystem accounts, both policy uses and spatial levels of interest (e.g., national, regional, local) can vary significantly, potentially leading to misalignments among expected policy uses at different scales. Pioneer practitioners in Australia^26^ highlight that the framing of urban ecosystem accounts is strongly shaped by initial policy purposes and the spatial level at which they are developed, which differ across implementations.

## 3. Discussion

Our work highlights a global shortage of consistent pilots on urban ecosystem accounting , which partly explains the lack of lessons and approaches (Table 2) for several challenges (Table 1). As illustrated in Fig.1, large regions, including most of Latin America, Asia, and Africa, appear to have neither tested SEEA-EA general accounts nor developed specific urban ecosystem accounts. This finding aligns with the gaps in SEEA-EA implementation identified in the UN’s 2023 Global Benchmark^157^. In Europe, Lange and colleagues observed that only a few countries have tested all five SEEA-EA general accounting tables, and thematic urban ecosystem accounts have rarely been tested^158^. This scarcity of pilots partially explains the absence of complete, consistent examples that could serve as practical guidance for ecosystem accounting implementations (Ch.20). Developing such examples for thematic urban ecosystem accounts will require first defining solutions for the challenges identified and implementing them based on a logical prioritization.

### 3.1. From approaches and lessons learned towards specific solutions

For the challenges related to the characterization and classification of urban ecosystems, we identified approaches or lessons focused on specific aspects rather than comprehensive solutions.

Despite the lack of scientific consensus on which assets belong to urban ecosystems (Ch.1), it is increasingly clear to scholars that not all artificial surfaces should be considered urban. This perspective is also gaining recognition among policymakers, as illustrated by a recent amendment to the regulation on environmental accounts in the European Union^159^. As required by SEEA-EA general accounts, the amendment differentiates a broad set of mutually exclusive ecosystem types, including “Settlements and Other Artificial Areas”, which represent artificial surfaces. The amendment also refers to urban ecosystems, named “cities and their adjacent towns and suburbs”, explicitly recognizing that urban ecosystems comprise multiple types of ecosystem assets, not just artificial surfaces. This growing differentiation between urban ecosystems and artificial surfaces, reinforced by the EU policymaking example, points to two partial solutions (as illustrated on the top of Fig.3 – Solution [Sol.] 1): i) using a term other than “urban ecosystems” when referring solely to artificial surfaces, as not all belong to urban ecosystems; and ii) compiling urban accounts exclusively through thematic ecosystem accounting to avoid confusion, breaking mutual exclusivity, and reporting incomplete representations of urban ecosystems.

**Fig. 3.**
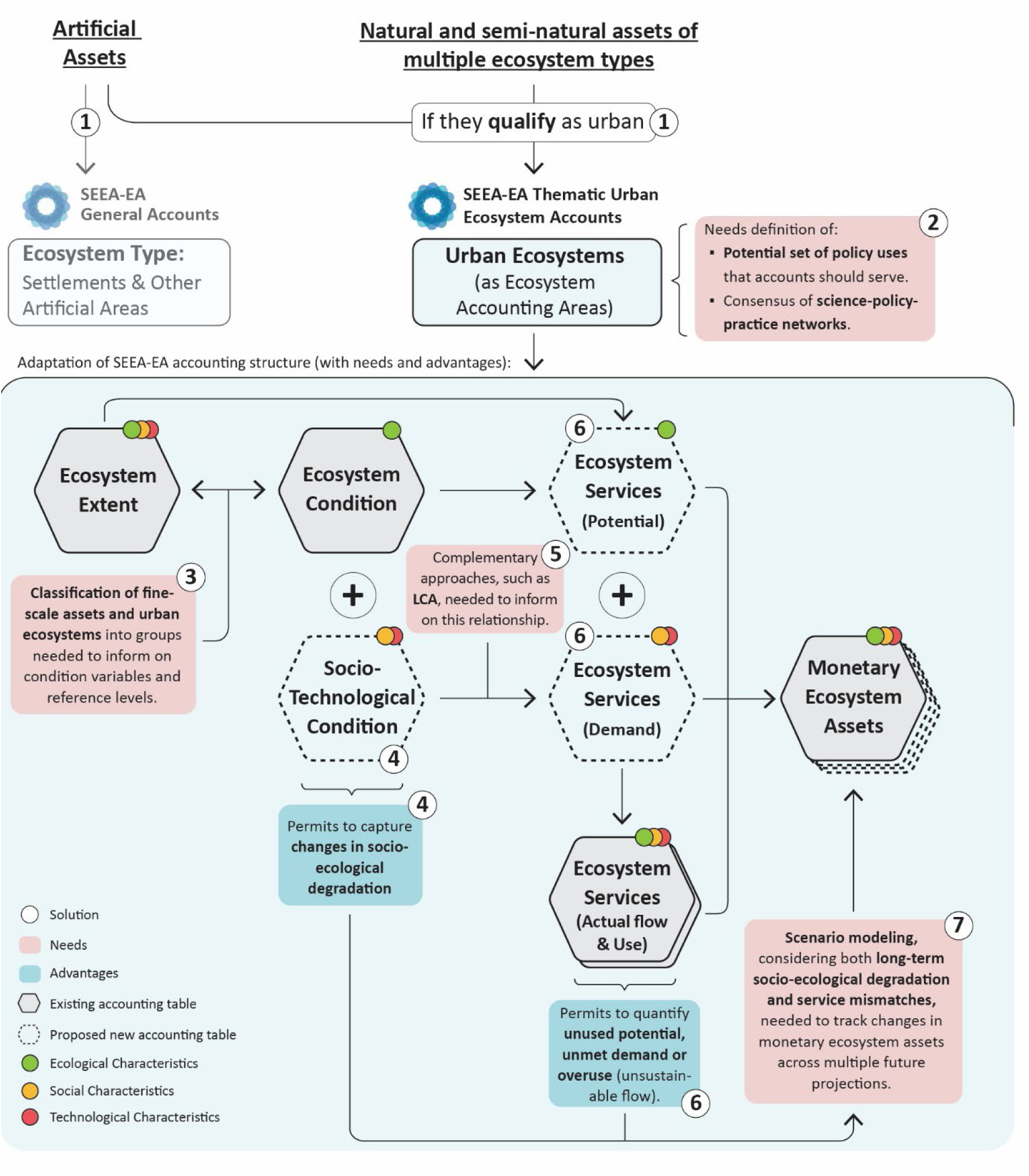
Visual summary of key solutions based on existing lessons and approaches. The figure presents solutions in the form of: decisions, actions to be implemented at specific steps (needs), and new accounting tables to be integrated into the SEEA-EA structure for thematic urban ecosystem accounting. For each proposed accounting table, the visual summary highlights its advantages, as challenges it may help to address, arising from the combined use of existing and newly proposed accounting tables. Additionally, the figure indicates whether ecological, social, and technological characteristics are relevant for defining each accounting table or estimating their values.

Unlike the previous challenge, the delineation of urban ecosystems (Ch.2), lacks established clear approaches from both scholarly or policymaking perspectives. This is illustrated in the EU Nature Restoration Regulation^160^, where three interrelated approaches for delineating urban ecosystem areas are proposed: i) selecting (urban) municipalities based on the degree of urbanization; ii) restricting the boundaries to highly populated zones of urban municipalities; and iii) expanding the restricted boundaries to include a peri-urban buffer. While delineating ecosystem accounting areas is also difficult in other mosaic-like ecosystems, such as wetlands, and in ecosystems with highly variable ecotones, like coastal ecosystems, this challenge partly differs for urban ecosystems. In other ecosystems, asset characterization and boundary delineation are typically conducted by experts and generally accepted. In contrast, characterizing (Ch.1) and delineating (Ch.2) urban ecosystems must be credible and legitimate not only scientifically but also to other stakeholders, including policymakers, adding constraints to the development of a coherent framework for thematic urban ecosystem accounts (Fig.3 – Solution 2).

For challenges related to tracking socio-ecological degradation, impacts and dependencies of urban ecosystems, approaches and univocal lessons point towards clear solutions.

Integrating social and technological variables into condition accounts would enable a more comprehensive assessment of urban degradation (Ch.5). However, this may conflict with rules of SEEA-EA general accounts, where ecosystem condition focuses on abiotic and biotic characteristics, strict ecological degradation, and there is a widely agreed-upon typology of condition variables. Directly modifying condition accounts in thematic urban ecosystem accounts could impact the interoperability with SEEA-EA general accounts. As a potential solution, we propose to create a “sister” condition table, specific to thematic accounts, in which only socio-technological characteristics are represented (Fig.3 – Sol.4). This new table, whose structure must be agreed upon, combined with the traditional condition table, would help to capture socio-ecological degradation in urban ecosystems more comprehensively. Additionally, replacing ecological integrity with ecological health or urban ecosystem health as a conceptual basis for defining reference conditions in urban ecosystems (Ch.6) could better capture changes in socio-ecological-technological conditions, though these new concepts require refinement and testing for ecosystem accounting purposes. Furthermore, classifying urban ecosystems, and their fine-scale assets, into coherent groups would enable the tailoring of condition variables and reference conditions to specific groups (Fig.4 – Sol. 2), thereby, informing more accurately about urban degradation.

**Fig. 4.**
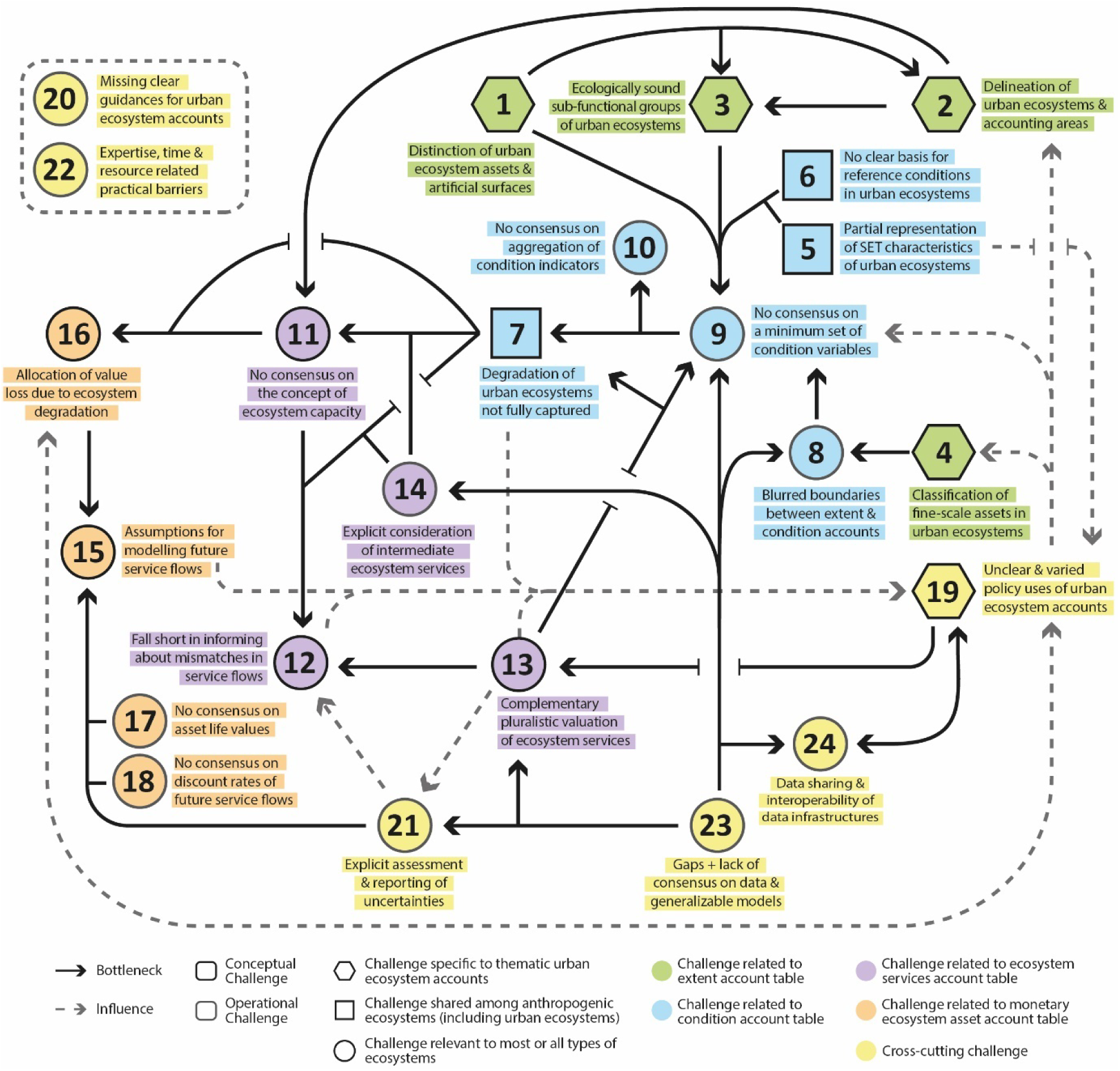
Interrelations among the identified challenges. Challenges are represented by their Ids (as used in Table 1), along with brief descriptive labels. The colours indicate the type of ecosystem account each challenge pertains to. Black arrows denote challenges (arrowheads) that are constrained by specific bottlenecks (end of arrow), while grey dashed arrows indicate challenges (arrowheads) that are influenced by others (end of arrow) but do not act as strict bottlenecks. The two operational challenges inside the dashed rectangle are indirectly influenced by how the remaining challenges are resolved. This network of interrelations help determine which challenges need to be addressed first, as well as those that should be tackled simultaneously and in a coordinated manner. Further details on specific interrelations are provided in the Supplementary Information (SI5).

In addition to the above solutions, integrating complementary natural capital approaches with a strong focus on the technosphere, such as life-cycle assessment, would help capture how social and technological characteristics shape services demand (Fig.3 – Sol.5). This integration would contribute to identifying mismatches in ecosystem service flows (Ch.12), i.e., unused potential, unmet demand, and overuse (unsustainable service flow), thus capturing the socio-ecological impacts and dependencies of urban ecosystems more accurately. As stressed in the literature^58,97^, addressing Ch.12 first requires explicitly reporting ecosystem service potential and ecosystem service demand as individual mandatory accounting tables (Fig.3 – Sol.6). This is not a current requisite under SEEA-EA general ecosystem accounts, which consider these accounting tables auxiliary.

The integration of scenario modelling into monetary ecosystem asset accounts would also enable tracking impacts on assets’ value across different future projections while making explicit the assumptions (Ch.15) used in each scenario (Fig.3 – Sol.7). Ideally, these scenarios should consider the effects of socio-ecological degradation and long-term service mismatches, leveraging the solutions proposed for condition and service accounts. Additionally, scenarios should integrate social and technological variables alongside climatic and environmental factors to provide more realistic projections. Scenario modelling further underscores the need for explicitly accounting of uncertainty (Ch.21), including propagated uncertainty, which can initially be addressed through simple techniques, as suggested in the literature^26^.

While the proposed scheme may help address some of the open issues, solutions for others remain problematic. For certain challenges, such as defining a clear set of policy issues (Ch.19), no single correct solution exists, and resolution depends not only on scientific consensus but also on a broad agreement across science-policy-practice networks. Lessons proposed for Ch.19, 22 and 24 lead us to key recommendations for achieving this consensus: i) focus on a limited set of policy purposes, to prevent ecosystem accounting from being perceived as a ‘one-size-fits-all’ tool, which could diminish its effectiveness; ii) engage key stakeholders early in the process, ensuring representation from both producers and end users with diverse expertise; iii) foster regional networks and communities of practice to help identify policy needs across different spatial levels and regional contexts.

### 3.2. Interrelations of challenges and priorities for actions

As illustrated in Fig.4, challenges are closely interconnected. These connections can either be bottlenecks, where one challenge hinders the resolution of another, or influences, shaping how issues are addressed or determining the most suitable approaches when no single solution exists. Therefore, it is important not only to derive solutions to individual challenges but also to consider their interconnections to establish a logical prioritization for resolution.

Fig.4 clarifies that some challenges depend on resolving multiple levels of interrelated dependencies. This is particularly true for challenges related to ecosystem service and monetary ecosystem asset accounts. For example, several interconnected challenges must be addressed before resolving shortcomings in ecosystem service accounts for capturing mismatches in service flows (Ch.12). Among other aspects, this requires consensus on what constitutes ecosystem capacity, how it should be assessed (e.g., service-by-service or systemically), and when it is exceeded (Ch.11). Without these steps, it is impossible to capture overuse (unsustainable service flows), as this requires an unequivocal definition of ecosystem capacity. In turn, Ch.11 itself depends on resolving bottlenecks related to ecosystem extent and condition (Fig. 4). Thus, challenges must be tackled with a logical prioritization, avoiding premature attempts to address those constrained by multiple bottlenecks, which would be counterproductive and may necessitate repeated revisions.

Instead, certain challenges related to extent (Ch.1-4), cross-cutting (Ch.19,21,23,24), and condition (Ch.5,6) are primary bottlenecks, being minimally or not constrained by other challenges. Prioritizing their resolution will unlock progress for other challenges. For instance, defining the policy uses that accounts should serve (Ch.19) strongly influences the development of a consistent framework for thematic urban ecosystem accounts, as policy needs strongly shape implementation^26^ and influence how multiple other challenges are addressed. Thus, to achieve a coherent global framework for thematic urban ecosystem accounting, it is a top priority to define early on a clear and consistent set of potential policy uses, agreed at international level.

Addressing primary bottlenecks that are not urban-specific will also benefit SEEA-EA general accounting and other ecosystem types. However, resolving these challenges cannot emerge solely from within the urban ecosystem accounting community, requiring broader consensus to ensure interoperability with SEEA-EA general accounts. Consequently, experts in thematic urban ecosystem accounting should first focus on challenges specific to urban or anthropogenic ecosystems, and minimally constrained (Ch.1–6,19), as these require less coordination. Meanwhile, cross-cutting and condition challenges relevant to most ecosystem types and similarly unconstrained (Ch.8,20–24) should be addressed next, in collaboration with SEEA-EA general experts. SI-5 offers further details on specific interrelations, including those not discussed here.

Addressing the identified challenges through a stepwise approach, building on existing lessons and approaches, will accelerate the advancement of standardized thematic urban ecosystem accounts. These accounts may become crucial for enhancing our understanding of urban ecosystems and supporting their sustainable management in a globally coherent manner. However, even once these challenges are resolved, ecosystem accounting, like any framework, will still have inherent limitations. While it can serve as a valuable tool for informing policies that promote urban sustainability, it should not be regarded as the sole tool.

## 4. Methods

Challenges, lessons, and approaches for urban ecosystem accounts were identified through a structured review of scientific and grey literature, following three consecutive steps.

First, key SEEA-EA documents^13,25,161^ were reviewed. Second, a systematic review of scientific and grey literature on ecosystem accounting was conducted. Third, a thematic review of additional scientific literature (not-accounting related) was carried out for those urban-specific challenges that go beyond accounting, i.e., remaining open issues in urban sciences. Each step is detailed below and visually summarised in SI2. The raw data gathered in the review can be consulted in SI6.

### 4.1. Step 1: Review of key SEEA-EA documents

The review began with the SEEA-EA Handbook^13^, including its section on a research and development agenda. It then extended to pitches (presentations) from the Virtual Expert Forum of SEEA experimental ecosystem accounting^161^ and discussion papers of the working groups for the SEEA experimental ecosystem accounting revision, available on the SEEA website^25^. Some of these discussion papers had already been published as peer-review articles or technical reports, or referenced additional relevant literature. This step aimed to compile an initial list of key challenges, drawing from both the SEEA-EA Handbook and broader expert discussions in the Forum and Working Groups, capturing details that may not have been included in the handbook.

### 4.2. Step 2: Systematic review of ecosystem accounting literature

The review combined an iterative literature review via an online “literature mapping tool” with a traditional systematic literature review.

In the iterative approach, the key SEEA-EA documents and references of interest cited in them were used as input data to identify other studies by making use of the online tool Connected Papers (https://connectedpapers.com). This tool uses an input document (e.g., peer-reviewed paper) to build a graph of documents (e.g., papers, books, chapters) that share references and co-citations with the input^162–164^. This approach helps to overcome typical issues of systematic reviews (e.g., missing keywords in the search strings) that could lead to missing key references^163^. It might help to find relevant references in emergent areas of studies where diverse keywords are still in use, as could be the case in urban ecosystem accounting studies. Prior and derived works were screened as detailed below and once new relevant studies were identified, they were also used as an input in Connected Papers. By iterating the process until no new relevant paper was appearing, a comprehensive list of studies was retrieved

The traditional systematic review was applied to ensure completeness, helping to identify additional literature. It followed a traditional systematic review approach, similar to previous studies^165,166^, and consistently with PRISMA guidelines^167,168^. The literature was gathered from Web of Science, Scopus, Google Scholar, and Google. In all cases, the broad search string “ecosystem account*” was used and the studies were limited to those written in English and published after 2012. This is the year in which the SEEA experimental ecosystem accounting framework, that led to the SEEA-EA, was developed^169^. The search string was intentionally broad to avoid missing documents of potential relevance. The keyword “urban” was intentionally excluded since there might be studies that integrate accounts for more than one ecosystem, without explicitly referring to “urban” (or use a related word) in the title, abstract or keywords. There might be also cases focused on other ecosystems or type of thematic ecosystem accounts, such as ocean, that include also challenges of relevance for thematic urban ecosystem accounts. The traditional approach helped to make the systematic review more exhaustive ensuring that relevant references were not missed. Moreover, the inclusion of Google Scholar and Google was useful to identify technical grey literature of potential interest, which was not captured in Web of Science or Scopus.

In the traditional systematic review, first, the title, abstracts and keywords of the studies gathered were screened retaining only: i) urban ecosystem accounts and related studies; ii) broad ecosystem accounts that included urban ecosystems; iii) other thematic ecosystem accounts, which discussed challenges that might be also relevant for thematic urban ecosystem accounts. The same criteria were then applied to further screen the full text of retained papers. Only studies applying the SEEA-EA framework, or its previous experimental version, were retained. As an exception, retained publications also included: i) studies discussing general shortcomings of ecosystem accounting frameworks, which apply also to SEEA-EA; ii) studies comparing advantages and disadvantages of SEEA-EA against other frameworks, provided that the challenges discussed were relevant for thematic urban ecosystem accounts. Studies that introduce concepts or methodologies related to SEEA-EA without specifically addressing conceptual/operational challenges for urban ecosystems or presenting lessons learnt were discarded. The same screening procedure and criteria were used for the studies gathered through the iterative approach. In that case, the screening of the abstract and full text was done before each new iteration, since retained papers were those used as inputs in new iterations.

The traditional systematic review and iterative literature review helped to identify challenges missed in key SEEA-EA documents as well as to further inform and better organise challenges already included in the initial draft list. After analysing all the studies, the challenges were organised in a final list, each of them corresponding to an aggregation of specific problems around a common theme. Challenges were classified in conceptual or operational ones, and according to: i) the type of ecosystem accounting table to which they refer; and ii) their urban specificity. Conceptual challenges refer to the development or refinement of theoretical concepts onto which the implementation of thematic urban ecosystem accounts should be based. Operational challenges refer to practical developments needed to set up clear and coherent procedures for implementing thematic urban ecosystem accounts. Based on the type of accounting table to which they referred, challenges were classified as: i) ecosystem extent, ii) ecosystem condition, iii) ecosystem services (biophysical and monetary), or iv) monetary ecosystem asset. Challenges relevant to more than one type of accounting table were classified as cross-cutting. Based on the urban specificity, challenges were classified as: i) specific to urban ecosystem accounts; ii) shared among accounts for anthropogenic ecosystems; iii) relevant to accounts of most or all types of ecosystems, including urban ecosystems.

The traditional systematic review and iterative literature review also helped to identify additional publications from which lessons and approaches could be drawn. Lessons are outcomes that instruct univocally on solutions to address a challenge or on relevant aspects to consider. Instead, approaches showcase alternative solutions followed by different authors for structuring an aspect of thematic urban ecosystem accounts, but which are not solved univocally among authors. Then the lack of an agreement might lead to diverse (incompatible) accounting structures, unless consensus is reached. The aggregation and organisation of lessons and approaches considered the specific challenge(s) to which they were related.

Besides recording challenges and lessons, the following data were recorded for each selected document: i) the type of document (scientific or grey literature); ii) the presence of a case study and its country; iii) the ecosystem types covered by the accounting (urban, multiple ecosystems including urban, other); iv) the focus of the ecosystem account (specific components or the entire ecosystem); and v) the type of ecosystem accounts (extent, condition, services in biophysical units [potential, actual flow - supply, actual flow - use], services in monetary units [potential, actual flow – supply, actual flow – use], and monetary ecosystem assets). These data provided contextual information about the retained ecosystem accounting studies.

### 4.3. Step 3: Thematic review of non-accounting literature

A thematic review of additional scientific literature (non-accounting related) was conducted to complement the collection of lessons and approaches for urban specific challenges that go beyond accounting. Specifically, the thematic review focused on challenges related to classification of urban ecosystems, suitable theoretical references for urban ecosystem condition or status, and appropriate evaluation of ecological degradation in urban ecosystems. These challenges, and associated lessons and approaches, belong to “hot topics” in urban sciences that have been already discussed by academics also outside ecosystem accounting. Therefore, in those cases, scientific studies outside ecosystem accounting were considered valuable sources for lessons and approaches.

The thematic review replicated the procedure of the iterative systematic literature review. Initial input documents consisted of studies identified in the systematic review, supplemented by additional key papers known to the authors.

## Supplementary Information

- **Supplementary Information 1.** Glossary of key ecosystem accounting terminology.
- **Supplementary Information 2.** Methodological workflow and overview of the data gathered in the literature review.
- **Supplementary Information 3.** Description of challenges.
- **Supplementary Information 4.** Description of lessons and approaches.
- **Supplementary Information 5**. Justification of the interrelations among challenges.
- **Supplementary Information 6.** Data and metadata from documents in the structured review of scientific and grey literature on urban ecosystem accounts.

## Acknowledgements

The authors used ChatGPT-4 to assist with proofreading and writing adjustments. All content was written, reviewed, and edited by the authors. All data analysis, review, and editing were conducted solely by the authors without the use of AI tools. JBA and RC acknowledge support from the National Biodiversity Future Centre (NBFC) project funded by the European Union’s NextGenerationEU, National Recovery and Resilience Plan (NRRP), CN00000033, CUP, H43C22000530001. Details of further funding sources supporting this research will be included at a later stage.

