## Supplementary Information 1 for "Advancing the global statistical standard for urban ecosystem accounts"

**Supplementary Information 1: Glossary of key ecosystem accounting terminology.**

The definitions and explanations of terms and terminology presented here are drawn from the SEEA-EA book[1], the EU-wide methodology report of the Joint Research Centre European Commission[2], and the INCA Platform of the Joint Research Centre of the European Commission[3]. They are provided to assist readers who may not be entirely familiar with ecosystem accounting in understanding the basic terminology. For more comprehensive explanations or additional details, please refer to the SEEA-EA framework. Terms that are still under discussion, such as 'ecosystem capacity,' have been excluded even when they are included in the glossaries indicated above.

**Ecosystem accounting area:** is the geographical territory for which an ecosystem account is compiled (extracted from the SEEA-EA Glossary).

**Ecosystem assets:** are contiguous space of a specific ecosystem type characterized by a distinct set of biotic and abiotic components and their interactions (extracted from the SEEA-EA Glossary).

**Ecosystem asset life:** is the time over which an ecosystem asset is expected to generate ecosystem services (extracted from the SEEA-EA Glossary).

**Ecosystem condition characteristics:** are the system properties of the ecosystem and its major abiotic and biotic components (water, soil, topography, vegetation, biomass, habitat and species) with examples of characteristics including vegetation type, water quality and soil type (extracted from the SEEA-EA Glossary).

**Ecosystem condition typology:** is a hierarchical typology for organizing data on ecosystem condition characteristics (extracted from the SEEA-EA Glossary).

**Ecosystem condition variables:** are quantitative metrics describing individual characteristics of an ecosystem asset (extracted from the SEEA-EA Glossary).

---

[1] https://seea.un.org/sites/seea.un.org/files/documents/EA/seea_ea_f124_web_12dec24.pdf
[2] https://op.europa.eu/en/publication-detail/-/publication/912e03a9-3fac-11ed-92ed-01aa75ed71a1/language-en
[3] https://ecosystem-accounts.jrc.ec.europa.eu/glossary

**Ecosystem degradation:** is the decrease in the value of an ecosystem asset over an accounting period that is associated with a decline in the condition of an ecosystem asset during that accounting period (extracted from the SEEA-EA Glossary).

**Ecosystem extent:** is the size of an ecosystem asset (extracted from the SEEA-EA Glossary).

**Ecosystem service demand:** it represents the need for specific ecosystem services by society, particular stakeholder groups or individuals. This assessment is not for filling Supply and Use table, but complementary accounting tables (Extracted from the Glossary of the INCA Platform of the Joint Research Centre).

**Ecosystem service potential:** Ecosystem contribution irrespective whether there is an ecosystem service demand or not. It measures and map the supply from the ecosystem side that eventually becomes actual flow/use once it interacts with the ecosystem service demand. This assessment is not for filling supply and use table, but complementary accounting tables. (Extracted from the Glossary of the INCA Platform of the Joint Research Centre).

**Ecosystem service use (actual flow):** The match between ecosystem service potential and ecosystem service demand that corresponds to the use of ecosystem service flows. This is the assessment that fills the ecosystem service supply and use tables. The actual flow is assessed in both physical and monetary terms (Extracted from the Glossary of the INCA Platform of the Joint Research Centre).

**Ecosystem type:** reflects a distinct set of abiotic and biotic components and their interactions (extracted from the SEEA-EA Glossary).

**Homogeneous ecosystem areas:** A detailed ecosystem classification relevant for the assessment of ecosystem condition that splits ecosystem types into more homogeneous areas regarding factors such as land cover data, climate, landform, human presence. This permits to split an ecosystem type into sub-groups of reduced variability, helping to define meaningful reference levels for ecosystem condition variables (Explanation based on content from the Section 3.3 of the EU-Wide Methodology Report).

**Reference condition:** is the condition against which past, present and future ecosystem condition is compared to in order to measure relative change over time (extracted from the SEEA-EA Glossary).

**Reference levels:** is the value of a variable at the reference condition, against which it is meaningful to compare past, present or future measured values of the variable (extracted from the SEEA-EA Glossary).
