## Supplementary Information 2 for "Advancing the global statistical standard for urban ecosystem accounts"

**Supplementary Information 2: Methodological workflow and overview of the data gathered in the literature review.**

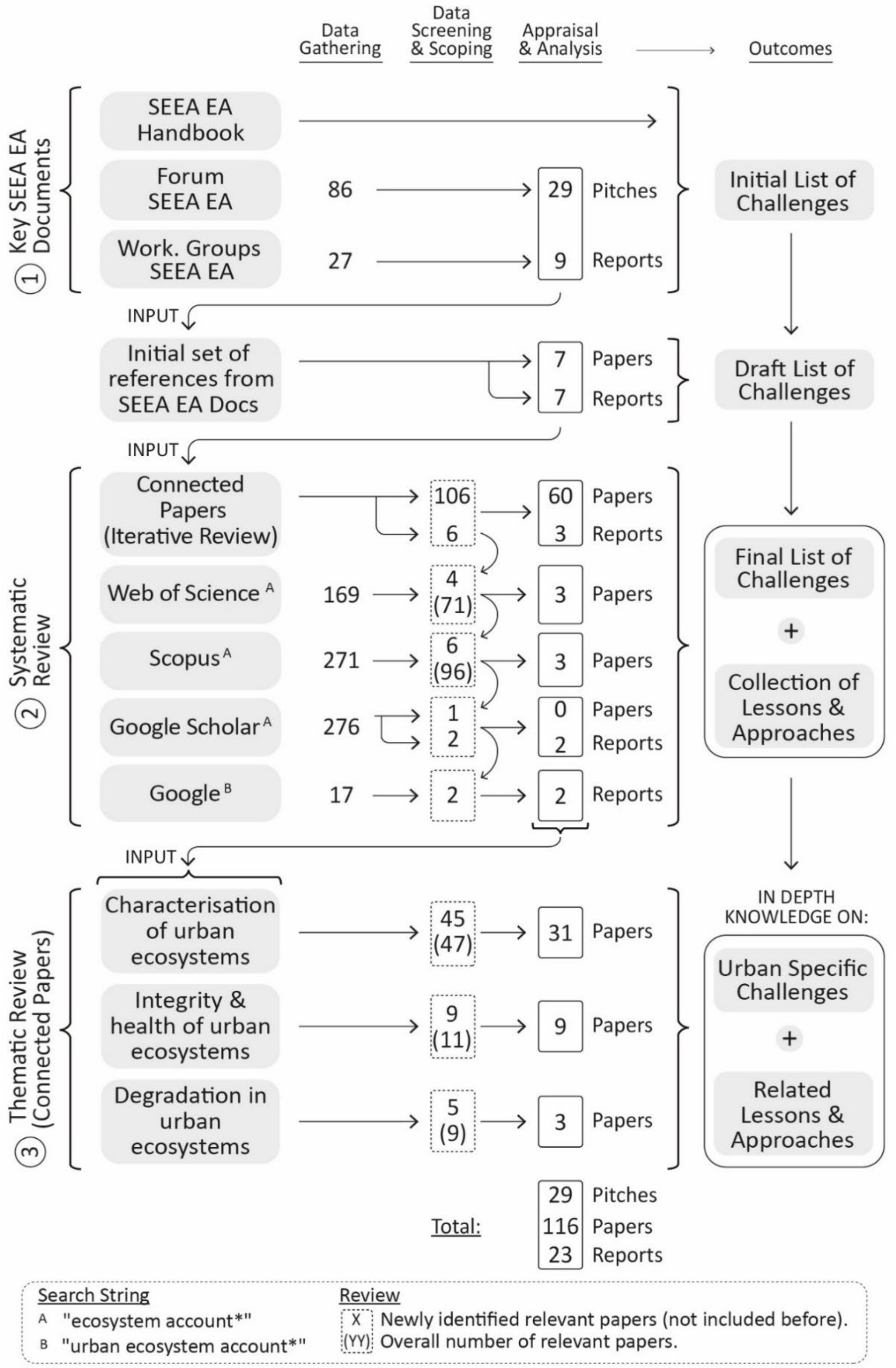


**Figure SI.2.1** |Representation of the methodological workflow of the three consecutive parts of the literature review and number of papers gathered and retained in each part. By pitches is understood the presentations in the Forum SEEA EA, which already inform on challenges.

**Overview of contextual data gathered in the literature review, in terms of:**

1. <u>Scope of accounting exercises and types of accounting tables considered in the set of studies</u>

In the review, 130 documents were retained, of which 87 were from SEEA EA documents and the grey and scientific literature on ecosystem accounting.

From those 87, 59% included practical or theoretical ecosystem accounting exercises. 39% of them were exercises with various ecosystems that included built areas, settlements, towns, cities, or urban ecosystems among the ecosystem types. 12% focused only on urban ecosystems, closer to SEEA EA thematic accounts than standard accounts. Aside from a few European-level exercises, ecosystem accounting was tested in only 19 countries at local, regional, or national levels (Fig. 1a). Only 6 countries had exercises specifically on urban ecosystem accounting (Fig. 1a).

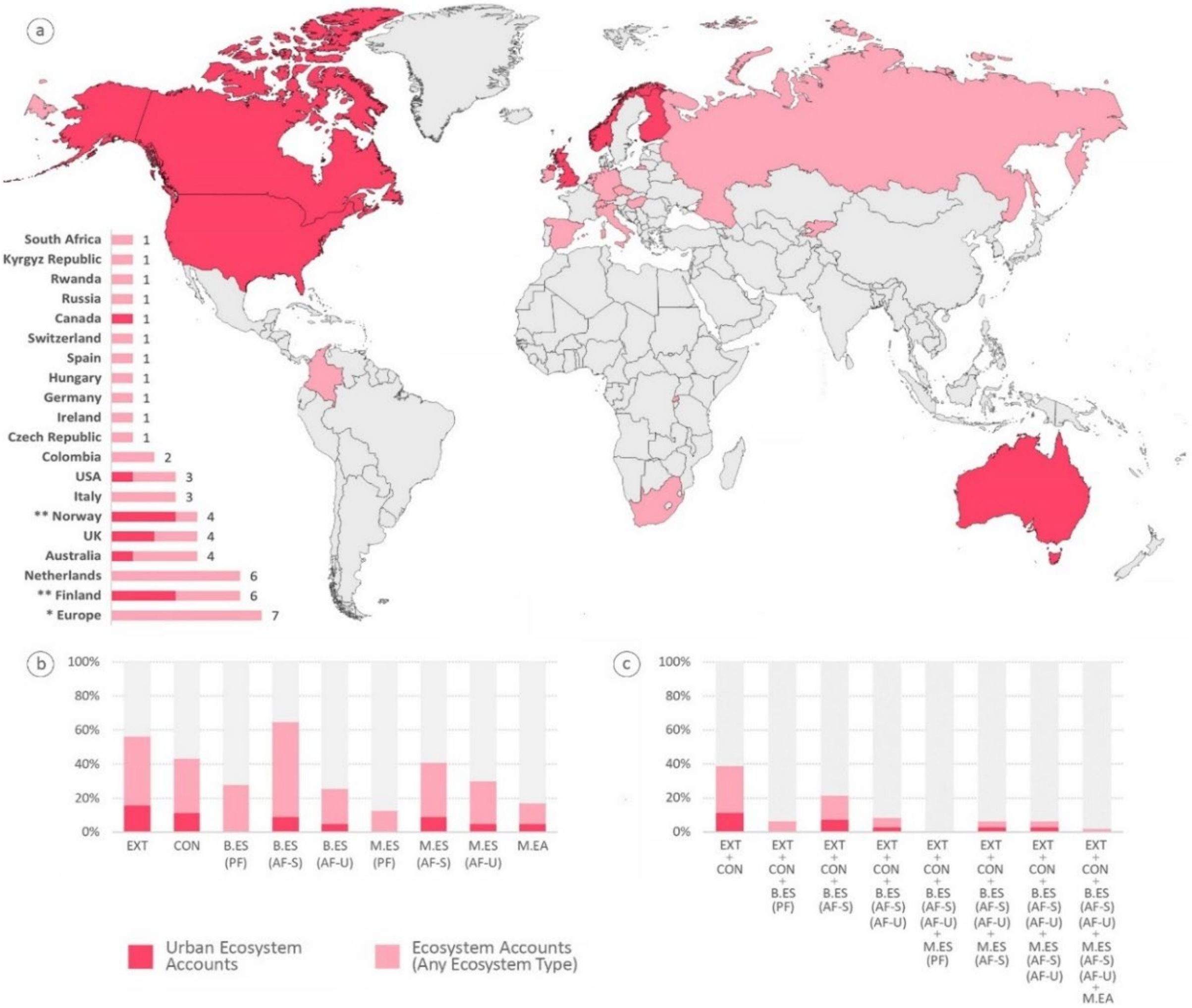


Figure SI.2.2 | Extended Figure 1 of case studies of ecosystem accounts (pink), distinguishing those focused only on urban ecosystems (red). **a)** World map showing countries where ecosystem accounts were tested in the literature reviewed. The bar chart displays the number of case studies per country. **b)** Share of documents, within ecosystem accounting exercises, that included specific types of accounts. **c)** Share of documents, within ecosystem accounting exercises, that included specific combinations of multiple types of accounts. **Note:** *Europe corresponds to an aggregation of ecosystem accounts developed at the European level, which differ in their boundaries. Some include EEA countries, EU-27, EU-28, or EU-28 with a few other European countries. **One document includes three case studies from Finland and one from Norway, which are counted as independent cases in Fig 1a). List of acronyms used: EXT = Ecosystem Extent Account; CON = Ecosystem Condition Account; B.ES (PF) = Biophysical Ecosystem Services Account (Potential Flow); B.ES (AF-S) = Biophysical Ecosystem Services Account (Actual Flow - Supply); B.ES (AF-U) = Biophysical Ecosystem Services Account (Actual Flow - Use); M.ES (PF) = Monetary Ecosystem Services Account (Potential Flow); M.ES (AF-S) = Monetary Ecosystem Services Account (Actual Flow – Supply); M.ES (AF-U) = Monetary Ecosystem Services Account (Actual Flow – Use); M.EA = Monetary Ecosystem Assets Account.

There is at least one example of each type of accounting table (e.g., extent, condition), although this is not the case for exercises focused on urban ecosystem accounts (Fig. 1b). No exercise included a complete set of all types of accounting tables (Fig. 1c). Ecosystems services potential flow accounts (biophysical and monetary) and monetary ecosystem asset accounts were among the least common in the exercises, urban-specific or not.

2. Weight of conceptual and operational challenges in the set of studies.

Challenges were identified in 23 key SEEA-EA documents and in 89 of the systematic and thematic reviews. Both, conceptual and operational challenges were frequently discussed in the literature. In the systematic and thematic reviews, approximately 76% of the 89 documents refer to conceptual challenges, 55% to operational, and 31% to both groups. In terms of conceptual challenges, 40% of the 89 documents focused only on those specific to urban ecosystems, 10% on those specific to any anthropogenic ecosystem, and 38% on challenges shared among most or all ecosystems, with several papers referring to multiple categories. When considering challenges per type of accounting tables, 54% of the 89 documents focused on cross-cutting challenges, while challenges related to extent and ecosystem services were each considered in around 30% of the documents. Challenges for ecosystem condition and monetary ecosystem assets were less frequently mentioned, each appearing in just 16% of the documents.

3. Weight of lessons and approaches per type of challenge in the set of studies.

There are only missing lessons or approaches for challenges 8, 10, 14, 17, 18 and 20 as reported in Table 1. For challenge 20 (lack of complete and consistent examples of urban ecosystems accounts that can serve as guidance), Figure 1 confirms the lack of examples for all the types of accounting tables (Figure 1b), and those that include a complete combination of accounting tables (Figure 1c)

Compared to challenges, many more documents were focused on lessons and approaches for conceptual challenges (90%), with only a 20% focused on lessons or approaches for operational challenges, and a 13% focused on both groups. A great part of the documents, around 84% included lessons or approaches for urban or anthropogenic specific challenges. In fact, half of the documents focused on lessons and approaches related to ecosystem extent challenges, being all urban specific (Table 1). Lessons and approaches related to ecosystem condition and crosscutting challenges were each included in around 20% of the documents. Instead, only few documents included lessons or approaches for ecosystem services (10%), and monetary ecosystem assets (4%).
