## Supplementary Information 3 for "Advancing the global statistical standard for urban ecosystem accounts"

### Supplementary Information 3: Description of challenges

This supplementary information provides a detailed description of all the challenges, based on the information collected during the various steps of the review. These challenges are organized and numbered as in Table 1 of the manuscript. Deliberately, the authors have limited their own interpretation when describing the challenges, ensuring that the collection and summary of information presented here remain differentiated from the discussion and proposals presented in the main manuscript. However, in some instances, brief notes or clarifications have been included where necessary. These additions, while reflecting the authors' interpretation of specific challenges, aim to enhance clarity and facilitate comprehension of certain issues.

#### <u>Ecosystem Extent</u>

1. **Challenge: There is still no univocal distinction between urban ecosystem assets and artificial surfaces.**

The classification and identification of urban ecosystem remain inconsistent, leading to confusion and potential misinterpretations.

The concept of urban ecosystems tends to be confused with the concept of artificial surfaces, assuming mining activities, excavation sites, airports, landfills, rural and remote roads, production yards, and other land assets not directly related to urban ecosystems are classified as urban ecosystem assets [1]. For instance, there are studies that include mining activities or landfills within urban ecosystems, without clear justification or explanation, suggesting that "urban" is treated as synonymous with artificial surfaces [2] [3]. In other studies, assets classified as "Mineral extraction sites" or "green open spaces" are considered to correspond to "dispersed urban areas" as specific ecosystem assets [4]. However, this semantic qualitative classification is misleading to other scientists and to the general society.

In the case of mineral extraction sites, it represents a clear example of an ecosystem extent accounting classification that would lead to inadequate interpretations of the impacts derived from low-density urban areas. Similarly, green open spaces can occur in dense and dispersed urban areas. They need to be accounted for in accounts within an adequate aggregation of classes that would be easy to understand by people using the data obtained from accounts and would not confuse them due to the terminology used. In fact, some of these land assets (e.g., mineral extraction sites) may be single objects surrounded by natural ecosystems, and assuming they represent urban ecosystems as defined by the IUCN classification is questionable [1].

In some studies, urban ecosystem assets are often identified as all the assets inside what is defined as the built environment. In those cases, urban ecosystem assets are all the assets inside boundaries delineated based on built-up extents or population density [5]. Other studies face the following dichotomy: urban ecosystem assets can also be considered to belong to a single ecosystem type or can alternatively be understood as belonging to multiple ecosystem types, depending on the specific asset [6]. A recent work advocates differentiating urban ecosystems from what has been called "Settlements and other artificial areas" [7]. The latter represents ecosystem assets where most of the human population lives, and which also include significant areas for synanthropic species associated with urban habitats. It includes urban, industrial, commercial, and transport areas, highly artificial green urban areas, mines, dumping, and construction sites [7]. It does not matter if these assets are single assets surrounded by natural assets or belong to a large human settlement. It does not intend to define what an urban asset is, but what constitutes an artificial asset, helping to differentiate both concepts.

Clear and standardized criteria are necessary to differentiate urban ecosystem assets from mere artificial surfaces, ensuring coherent and comparable thematic urban ecosystem accounts and avoiding confusion among scientists and policymakers.

2. **Challenge: There is still no univocal procedure for the spatial delineation of urban ecosystems and their accounting areas.**

The classification and spatial delineation of urban ecosystems is not straightforward, primarily due to the inherent variability of urban environments and differing perceptions of what constitutes an urban ecosystem and its boundaries. The SEEA EEA discusses ecosystem accounting in urban areas, emphasizing the challenge of defining the extent of urban ecosystems [8].

Urban ecosystems are often characterized as mosaic landscapes, making it difficult to classify them into specific ecosystem types[1]. At a broad, top-down driven scale, an ecosystem asset might be viewed as urban, but at a detailed scale it might be a wetland, lake, cropland or other type of ecosystem [9].

The absence of objective, clear rules for identifying urban ecosystems further complicates the establishment of accounting boundaries[1]. Defining these boundaries may involve considering factors such as: i) size thresholds (e.g., minimum population or built-up area) for including urban areas in accounts; ii) the extent of peri-urban areas to include (if any); and iii) delineation methods [5 10 9]. The lack of standardization in urban area delineation approaches—such as using functional urban areas (FUA), municipal limits, or built-up areas with buffers—adds to this challenge. For example, the FUA approach may not align with common perceptions of urban areas, municipal limits often fail to capture the full extent of urban ecosystems, and buffer-based methods may lack relevance for policymaking (Wang et al., 2019).

Moreover, there is no clear consensus on what urban ecosystem accounts should track, which directly affects the scale and scope of delineating and classifying specific assets. These choices are influenced by the types of policy questions that urban ecosystem accounts aim to address [9].

The definition of urban ecosystems also has a significant impact on how peri-urban areas and transboundary effects are treated. Some studies define urban boundaries ecologically, incorporating rural hinterlands because of their focus on biodiversity and habitat connectivity. Others adopt a socio-economic perspective, including only peri-urban forests and nearby areas that influence urban residents' well-being, property values, and the perceived value of urban green spaces as substitute sites [5].

For instance, the UK's urban extent is delineated using the Office for National Statistics (ONS) Built-Up Areas (2011) dataset, modified to include peripheral nature areas where high levels of human-nature interaction occur, as well as larger urban green spaces such as city parks and rivers [11]. In the UK National Ecosystem Assessment's habitat classifications, urban ecosystem extent encompasses built-up areas and gardens, excluding transport infrastructure outside urban and rural settlements and structures like quarries in rural areas [12]. The degree of urbanization (DEGURB) classification represents urban ecosystems as densely populated and intermediate-density areas [13]. Another example is the Urban Morphological Zone (UMZ) used by the European Environmental Agency, defined as 'a set of urban areas lying less than 200 meters apart.' The UMZ focuses on specific CORINE land cover classes without considering population distribution, functional economic links, or commuting flows. As UMZs are not used for data collection but only for aggregating existing information, they primarily represent 'built-up areas' [13].

Urban ecosystem assets are often defined by their proximity to the built environment rather than specific ecological features. For example, distinguishing between an "urban river" and a "non-urban river" requires defining clear rules. This is a critical, non-trivial, step when developing urban ecosystem accounts [14]. The fragmented nature of urban ecosystems—spanning broad types like freshwaters and forests to small-scale features like street trees and green roofs—further complicates this process [14]. Therefore, urban ecosystem accounting must accommodate varying levels of detail, from individual green assets (e.g., street trees, green roofs) to larger urban areas with shared characteristics that can be linked to ecosystem services [6].

Additionally, the lack of standardized urban ecosystem delineation complicates comparisons across scales and regions [15] [16]. Several scholars state that administrative boundaries, often used as reference data, fail to capture the spatial continuity of urban built-up areas, leading to inconsistencies in international and temporal analyses [16]. Despite progress, achieving a universally accepted method for urban delineation remains an ongoing challenge [16], at least for the purposes of urban ecosystem accounts.

Scholars in urban ecology state that developing a standard set of measures to quantify urban gradients and improving their integration across studies may be critical to advancing our understanding of urban ecosystems. Such measures would enhance the comparability of urbanization metrics globally and will contribute to the development of robust urban ecosystem accounts [17].

3. **Challenge: At a global level, urban ecosystems still lack an ecologically sound, agreed-upon classification of homogeneous ecosystem areas, along with the principles to develop it.**

Urban ecosystems, whether defined by administrative boundaries, built-up extent, functional area, or other criteria, comprise a set of diverse land covers and uses and may exhibit significant differences in structure within specific types of urban ecosystems [9]. However, there is no broadly agreed-upon division of urban ecosystems into distinct types or groups.

While the delineation of ecosystem assets should emphasize an ecological perspective, land cover alone is insufficient to characterize an urban ecosystem. This prompts the question of which additional characteristics, beyond land use, should be considered [18]. For characterizing and typifying urban ecosystems, a standardized set of measures to quantify gradients and provide an 'urbanization context' for studies is crucial. This requires clarity on the most informative combinations of demographic, physical, and quantitative landscape metrics, as well as on the interactions between classification system characteristics (e.g., scale, typology) and study outcomes [17].

Urban research faces challenges in the comparability of urbanization measures across regions and time periods [16]. This issue arises from the lack of an agreed-upon typification of urban ecosystems. Efforts to describe urban ecosystems and develop typologies range from qualitative approaches rooted in architecture and planning to quantitative methods leveraging modern geographical analysis [19]. Furthermore, comparative studies that analyze the internal morphological structures of cities using Earth Observation (EO) data are limited to a small number of cities, and the specifications of the required or used input data rarely allow, for various reasons, a global approach [20]. As such, studies analyzing the intra-urban structure of cities or conceptualized models of cities for global empirical comparison based on consistent datasets do not yet exist [20].

This challenge has been recognized by ecosystem accounting scholars, who acknowledge that urban ecosystems lack consensus on such clustering, and that socio-ecological classification efforts remain in their early stages [21]. They also highlight the importance of typifying or classifying urban ecosystems for other ecosystem accounting steps, such as defining reference levels of ecosystem condition [21].

4. **Challenge: Urban ecosystems still lack a common ecologically sound classification of the fine-scale assets (patches) that compose them.**

Globally, urban ecosystem assets are often grouped into a single homogeneous class, in contrast to vegetated landscapes, which are categorized into 13 or more distinct classes [22]. Just as deciduous forests differ from evergreen forests and savannahs from shrublands, a single "urban" or "built-up" class limits our ability to understand the unique assemblages of neighborhoods within urban areas and their varied, often uneven, impacts on human and environmental well-being [22].

The SEEA-EA handbook suggests classifying fine-scale assets in urban ecosystems under the landscape or individual approaches depending on specific purposes, but multiple approaches might lead to issues of comparability and harmonization among ecosystem accounts (United Nations, 2020).Defining sub-types of fine-scale assets may involve a variety of criteria, including land use, land cover, intensity of use, density, property ownership, or combinations thereof (Wang et al., 2019).

Some scholars state that this classification of fine-scale assets is necessary for conducting an adequate assessment of the ecosystem services generated by urban ecosystems at the landscape level, as it requires urban areas to be subdivided into smaller patches that maintain structural diversity while being regrouped within homogeneous areas [6]. Other scholars state that most urban ecosystem service (ES) maps rely on readily available land use and cover (LUC) datasets, which lack the spatial and thematic detail necessary to capture fine-scale components that support ES provision. Although novel urban landscape classifications are emerging, such studies remain rare, and systematic model validation against primary data is uncommon [23].

As a consequence of the lack of a common fine-scale asset classification, there is also a lack of standardized variables and common datasets across cities, impeding cross-city comparisons [24]. Comparative research on the relationships between urban structures and ecological functions requires classification systems with comparable structures and spatial scales tailored to the functions of interest [24]. Fine spatial resolutions are especially important due to the high diversity and rapid changes in land cover within urban areas. Standardized classifications must also include land use or cover types with consistent meanings across cities to avoid inaccuracies, such as assumptions about vegetation proportions in residential areas [24]. Moreover, understanding the link between spatial heterogeneity and ecological processes in urban systems demands the quantification of fine-scale heterogeneity using structural elements that influence ecological processes [25]. Integrating built and non-built components into these classifications is essential for advancing such analyses [25].

The functional classification of fine-scale assets would help elucidate social and ecological relationships in urban ecosystems [26]. Despite recent fine-scale land cover classifications enabling more nuanced analyses, significant challenges persist in integrating spatial structure and configuration to support scalable, reproducible studies of urban form and process [26]. These challenges include the need for independent measurements of fine-scale spatial variability in environmental and ecological variables, particularly in the often-overlooked vertical dimension and three-dimensional variability of urban landscapes [26]. Currently, population or population density is among the main criteria for characterizing urban assets and urban development. However, these measures are, in most cases, chosen because of their widespread availability across cities, not necessarily because they are the most relevant [22].

### Ecosystem Condition

#### 5. Ecosystem condition variables offer just a partial representation of socio-ecological-technological characteristics driving the functioning and quality of urban ecosystems.

Current approaches to urban ecosystem accounting do not consider socio-economic dimensions, despite management, ownership, and governance influence ecosystem function and condition[1].
Ecosystem condition accounts, as part of the SEEA-EA, contain aggregated statistical information about the abiotic and biotic quality of an ecosystem. Among the approaches suggested to assess ecosystem condition, SEEA-EA proposes the use of biophysical indicators assessed against pre-industrialization baseline values, which act as reference levels [27]. However, this approach may not align with Indigenous contexts where bio-cultural interactions with ecosystems have existed for millennia [27]. One way to address this challenge in indigenous contexts (urban or not) may be the development of indicators for the relative bio-cultural condition of ecosystem assets that would complement existing ecological condition metrics [27]. This serves as an example of the limitations of ecosystem condition variables in

terms of the partial representation of the functioning and quality of ecosystems, in which human management and humans have a significant influence.

The representation of spatial context in ecosystem services (ES) literature spans five domains - biophysical structures, natural structures and processes, built structures and processes, individuals and society, and maintenance and governance. These domains highlight the interplay of ecological and social structures and processes in urban environments [28]. Mediating mechanisms such as human management, mobilization, allocation-appropriation, and appreciation influence the capacity of ecosystems to provide services, the realization of these services, and their perceived value [28]. By modifying the capacity of the ecosystem to provide a service, human management alters ecosystem conditions, and the role of this management risks not being captured since social and technological dimensions are not considered. In fact, in some cases, such as with urban trees, human mediation has been documented as highly relevant in influencing urban tree ES, followed by built structures and processes [28].

Data selection for ecosystem condition accounts must be refined; however, data that do not meet the selection criteria could be still considered as ancillary data (e.g., pressures), as they also provide relevant information about the system [29].

In terms of approaches, hybrid monitoring approaches that incorporate both transferable and context-specific indicators, are essential for capturing the full extent of people-nature systems [30]. Indicators should not only assess biophysical conditions but also monitor spiritual connections to the land and the wellbeing of people, which are integral components of ecosystem interactions [30]. Furthermore, the SEEA accounting system currently lacks measures of wellbeing at individual or societal levels [30]. To fully reflect the benefits of nature - including those derived from ecosystem services and stewardship activities - SEEA accounting systems should also incorporate measures of the 'state' or 'condition' of the human component of the interlinked and inseparable people-nature system [30]. For accounts to provide useful information on the system, they need to reflect the whole system, not just a part [30].

**6. Challenge: There is no clear conceptual basis for defining reference conditions in urban ecosystems, being ecological integrity inadequate as a default concept for them.**

This challenge stems from the need to establish anthropogenic reference conditions, yet more theoretical explanation is required to ensure consistency across different contexts [1]. In fact, defining a reference condition in urban ecosystems appears to be as much a policy issue as a scientific one [1]. Condition variables can be used to track whether conditions are improving or deteriorating relative to a target benefit, avoiding the necessity of defining a specific reference level [1].

It may be more appropriate to use target levels as baselines to assess indicators associated with anthropocentric conditions [31]. The distance from these target levels would infer quality, which could benefit policy applications but necessitate careful consideration of the scientific objectivity of this process [31]. The SEEA-EA framework recommends defining reference condition levels based on the natural state to develop accounts devoid of value judgments and free from implicit policy goals. However, later interpreting those reference conditions as an ecological debt would require linking the costs of reaching these levels to societal willingness to pay for ecosystem maintenance and restoration [32]. Under this approach, ecological debt can be estimated based on societal standards revealed by policy goals [32]. Yet, this raises practical challenges, including how collective willingness to pay can emerge from dynamic political processes involving diverse communities and how scientific, political, and administrative spheres can collaborate on target setting and implementation across scales [32].

The definition of reference levels as good ecological status can serve as a boundary object to bridge different communities without requiring them to drastically alter their specific optimal references [32] Scientific arenas warn of environmental limits and the risks of crossing thresholds, but such limits often carry normative content and leave trade-offs to public discussion [32]. Political processes, in turn, elaborate environmental targets to ensure the legitimacy of collective choices, balancing economic, social, and environmental considerations [32].

Moreover, according to some scholars, biological (ecological) integrity represents one endpoint on a gradient of ecological conditions and may not always be a feasible or desirable reference or goal for all places [33]. For lands under intensive human use such as cities a more reasonable goal would be ecological health [33]. Managing for ecological health includes preventing land degradation for future use and beyond the site [33].

**7. Challenge: Ecosystem condition accounts cannot fully capture socio-ecological degradation and indirect degradation in urban ecosystems.**

Ecosystem condition accounts are limited in representing socio-ecological degradation (both direct and indirect) of urban ecosystems. For example, urban demographic and economic growth place global pressure on ecosystems, urban expansion degrades ecosystems adjacent to urban areas, and disparities in access to ecosystem services within urban settlements may lead to unhealthy living environments [9]. These key issues, highlighted in the Millennium Ecosystem Assessment, are not adequately captured by urban ecosystem condition variables, which provide only a partial representation of socio-ecological degradation within urban ecosystems and of the degradation driven by urban drivers elsewhere [9].

The inclusion of externalities or disservices in extended System of National Accounts (SNA) requires recognizing ecosystems—or agents acting on their behalf—as entities affected by economic activities, especially when externalities impact both current and future flows of ecosystem services. In the latter case, ecosystem asset degradation becomes a critical consideration [34]. Overuse of services by humans, where actual flows exceed potential flows, contributes to degradation by decreasing the ecosystem's capacity to provide services over time. Although this link is neither direct nor linear across time and space, it is important to measure and record these differences to accurately reflect ecological degradation and ecosystem asset depreciation [35]. Current ecosystem accounts also fail to include indicators for resilience or to consider the probabilities of sudden collapses in overexploited ecosystems [36], or in systems (e.g., urban ecosystems) that depend on those overexploited ecosystems.

To gain a better understanding of ecological degradation, ecosystem condition, extent, and services accounts must work together to provide a comprehensive perspective, as degradation can occur even under sustainable use patterns [37].

**8. Challenge: There are difficulties establishing boundaries between ecosystem condition and ecosystem extent accounts, particularly at fine levels of ecosystem classification.**

The SEEA EEA framework highlights issues in defining urban ecosystem extent and differentiating extent from condition [8]. In fact, the expert working groups transitioning from the SEEA-EEA to the SEEA-EA indicated that urban thematic accounts tend to have a more blended view of extent and condition when compared to ecosystem accounts more broadly[1]. This statement is also supported by other scholarly works, which indicate that in highly modified urban landscapes the practical distinction between ecosystem extent and condition is often blurred and may not be useful for ecosystem service accounting [38]. For instance, asset-based approaches, such as those applied in pilots in Oslo, consider all vegetation cover part of the urban ecosystem extent while interpreting vegetation structure, including tree canopy extent and type, as indicators of condition [38]. This highlights the interconnected nature of these accounts in urban ecosystem classification [38].

**9. Challenge: There is no consensus yet on a common minimum set of ecosystem condition variables and on their reference levels.**

This challenge arises from various factors, including the reliance of countries on existing, non-harmonized monitoring programs, which complicates the establishment of a common framework [1]. Furthermore, the selection of condition variables and the definition of reference levels are influenced by the type of values considered -such as anthropocentric, ecocentric, intrinsic, or instrumental- in ecosystem condition accounts, which necessitates a clear understanding of their implications (Keith et al., 2019).

The SEEA EA framework must accommodate diverse biophysical, geographical, and socio-economic contexts, including realms, biomes, climatic zones, land use, and data availability. Such diversity makes it difficult to recommend concrete lists of metrics that are universally applicable [29]. While harmonized lists of fit-for-purpose indicators are desirable, achieving this requires globally consistent data availability supported by harmonized monitoring programs [29]. Additionally, linking data, methods, and metrics to specific accounting contexts and ecosystem types is critical for coherent global operationalization [29]. Robust assessments must account for the various structures and functions of ecosystems and balance intrinsic values (ecosystem integrity unrelated to ES supply) and instrumental values (how ES supply depends on ecosystem condition), this makes no single set of indicators suitable for all purposes [39]. Oversimplification and reliance on one-size-fits-all indicators risk losing critical information [39].

Despite SEEA EA framework establishes a sequential connection between ecosystem condition accounts and ecosystem service (ES) accounts, it lacks practical guidance on linking these components [40]. Variables used in condition accounts can inform ES supply and use tables, yet operationalizing this connection remains a challenge [40]. Indicators must balance transferability and context specificity; while some may be widely applicable, others must reflect the unique characteristics of specific ecosystems [30]. Beyond biophysical indicators, measures of spiritual and cultural connections to land and the human condition (e.g., “feeling good”) are essential for capturing the full extent of the interlinked people-nature system [30].

Moreover, as a specificity, urban ecosystems require standardized measures to quantify gradients of urbanization globally [17]. It would facilitate the comparability an integration of data between studies, since the urbanization gradient will be considered in an equivalent form [17]. These measures should address key combinations of demographic, physical, and landscape metrics; consider the interaction of classification systems; evaluate redundancy or correlation between measures; and enable ecological interpretation [17].

**10. Challenge: There is still insufficient evidence and consensus on the appropriate procedure(s) to aggregate ecosystem condition indicators into indices.**

Indicator selection and aggregation must be underpinned by a clear conceptual framework [1]. Establishing approaches to aggregation is recognized as a critical consideration for accounting; however, it remains inherently challenging in ecology [1]. The use of additive methods for aggregating variables necessitates simplifying the aggregated scores, in part to ensure comparability and facilitate communication with stakeholders [39]. Creating such simplified scores requires defining thresholds that qualify condition based on all characteristics, including their interactions, for which there is a general lack of empirical evidence [39]. These thresholds, often based on expert decisions and considering the distribution of summed scores, may have limitations, including being arbitrary, being unsuitable for interpreting values outside the originally established range, and creating a false sense of consistency [39].

<u>Ecosystem Services</u>

**11. Challenge: There is no consensus about the concept of ecosystem capacity, its connection to extent and condition and to other related concepts (e.g., degradation, sustainability, and resilience).**

There is still an absence of a clear and measurable definition of ecosystem capacity within the context of ecosystem accounting [1], as has also been recognized in the research agenda of the SEEA-EA. Currently, there are three conflicting perspectives that frame ecosystem capacity: the current individual ecosystem services (ES) perspective, the future individual ES perspective, and the systemic perspective [1]. In the current individual perspective, ecosystem capacity is generally calculated for individual ecosystem services rather than for ecosystems as a whole, providing transparency in the assessment process and enabling the identification of services or ecosystem types under threat [41]. However, this approach lacks integration across services, delaying the understanding of synergies and trade-offs that need to be investigated with ex-post analysis [41].

The concept of ecosystem capacity builds on ecosystem functions, which are ecological properties underlying service supply, yet few studies have quantified these functions, and no systematic framework exists for their measurement [42]. Capacity is usually based on the measurement of potential flow, which links it to ecosystem condition accounts and facilitates systematic assessments of degradation and sustainability in natural capital [41]. This potential flow approach allows a connection with sustainability thresholds, which, while often kept constant for simplicity, ideally vary based on context and asset conditions [43].

The relationship between ecosystem characteristics and ecosystem service capacities (EC ~ ES relationships) remains poorly understood, and guidance is lacking on how to define and apply ecosystem capacity, especially across provisioning, regulating, and cultural services [29] [42]. Additionally, knowledge gaps persist in specific contexts, such as the relationship between ecosystem quality and recreational use [44]. Although ecosystem capacity accounts aim to link ecosystem extent, condition, and services—fundamental to the SEEA EA framework—they remain undeveloped, being part of the SEEA-EA research agenda, and limit their utility in policy and management [45].

To address these issues, three considerations have been proposed for arriving at more precise capacity definitions [42]: (1) capacity should reflect an ecosystem asset's ability to provide specific services sustainably over time; (2) while capacity pertains to individual services, ecosystems deliver multiple interlinked services and the capacities for each of these services are also interlinked; and (3) capacity should be quantifiable in physical and monetary terms, supporting the conceptualization of ecosystems as assets.

In addition to capacity, it may also be relevant to understand which services could be provided by an ecosystem if it was managed differently [42]. Some scholars define this as capability, referring to the ecosystem's ability to sustainably generate one ecosystem service under current conditions and type of use, and irrespective of potential impacts of increasing supply on the supply of other ecosystem services [42]. Capability requires there to be a demand, since increasing the supply of a specific service is only meaningful if there is a demand for that service [42].

For services with rival characteristics, capacity-flow balances are meaningful. They are also useful in the case of congestible services, which are services that may exhibit thresholds of use intensity that impact sustainability, requiring careful management to maintain benefits for all users [46]. To move forward, a systemic approach is needed to measure ecosystem capacity and its links with condition, degradation, enhancement, and resilience[47].

**12. Challenge: Ecosystem service accounts fall short in informing whether there are mismatches in service flows that represent sustainability issues (e.g., degradation), unmet societal needs, and/or missed flows.**

Ecosystem service accounts often fail to adequately capture the relationship between potential flows, actual flows, and sustainable flows, leading to challenges in addressing mismatches and ensuring sustainable use. Potential flow, defined as the maximum flow an ecosystem can generate, differs from actual flow, which reflects what is currently utilized, and sustainable flow, which does not exceed regeneration rates [48]. However, the distinction between these flows is rarely assessed in a cohesive manner, with the complexity of integrating ecological and socio-economic systems complicating the quantification of actual flows [49]. For provisioning services, it is possible to quantify the difference between potential and sustainable flows [48]. For regulating and maintenance services it is possible to measure the sustainable flow once a sustainability threshold has been identified, but it is unclear whether it would be possible to measure potential flow [48]. This is a key point that needs to be addressed in order to make the accounting for ecosystem services operational and rigorous. Even establishing a sustainability threshold is not trivial because the conditions and vulnerability of ecosystems vary in space and time [48]. Moreover, the SEEA EA framework lacks components to directly address ecosystem potential, missing a vital link between condition accounts and ecosystem service tables, which could be pivotal for policy decisions [45].

The inability to record and analyze mismatches—where actual flow exceeds potential or sustainable flow—has critical implications for ecosystem degradation and long-term sustainability. Such mismatches indicate overuse that can lead to diminished future capacity to provide services [41]. These gaps are compounded by limitations in recording sustainable flows within frameworks such as SEEA-EEA, which do not yet adequately differentiate between sustainable and unsustainable ecosystem use, making it difficult to detect degradation risks [50]. This is a problem that persisted in SEEA-EA. Mapping both ecosystem service flows and ecosystem capacity could identify areas where flows exceed capacities, signaling unsustainable use and the need for intervention [51].

Additionally, the absence of an alert mechanism within current ecosystem service accounting frameworks perpetuates unsustainable practices. For example, computing Net Present Value (NPV) based on unsustainable flows (e.g., overused ES) can mislead policymakers by inflating perceived ecosystem capacity and reducing incentives for adopting sustainable practices [40]. This disconnection undermines efforts to manage ecosystems responsibly, as overuse may only become evident after ecosystems have degraded [40]. Furthermore, there are vulnerabilities associated with unmet demand for ecosystem services, which result from deficits in potential supply, while unused ecosystem service potential may remain harmless [40]. Another example of why is relevant to inform about mismatches in ecosystem service flows.

Finally, the separation between the benefits received from ecosystems and the actual services provided complicates efforts to address degradation. Without identifying causal relationships between enabling actors, beneficiaries, and service flows, policymakers lack the tools to plan and implement targeted environmental measures [52]. To resolve these challenges, it is imperative to enhance ecosystem service accounts with information on sustainable and potential flows, develop mismatch accounts, and integrate these into policy frameworks to safeguard ecosystem sustainability [35 52].

**13. Challenge: There is still missing a complementary pluralistic valuation of ecosystem services (and assets) that represents multiple systems of values.**

This challenge extends to the valuation of ecosystems and their services, as ecosystem accounting often disregards multiple valuation perspectives, raising questions about how to enhance or modify ecosystem accounting to better reflect a broad framing of values, as seen in IPBES frameworks[1]. Current valuation methods, aligned with the System of National Accounts (SNA), are limited in scope, excluding aspects like consumer surplus and welfare generation, thus failing to capture the total value of ecosystems [29 35]. Exchange prices for ecosystem services, reflecting current pricing mechanisms and market conditions, do not reflect the welfare generated by using natural capital, and therefore should not be interpreted as the total value of nature [35]. In fact, in ecosystem accounting monetary valuations do not address the contribution of ecosystem services to ecological or social production functions (ecological integrity and long-term well-being), with some authors arguing that this valuation must also include ecological and socio-cultural values that cannot be expressed in monetary terms [53].

Some authors have suggested breaching the concept of "nature as a service" by explicitly incorporating non-instrumental values into decisions that also represent spiritual and heritage values, cultural identity, and social cohesion, which are now invisible in decisions based on valuation outputs [53]. Indigenous perspectives are particularly underrepresented, despite their crucial role in ensuring culturally appropriate and equitable quantification of ecosystem services [30]. Indigenous-led approaches are essential to avoid the marginalization of their valuation systems in environmental and economic reporting and to align human–nature relationships with existing frameworks like the SEEA-EA [2754 30]. Relational values, central to many Indigenous worldviews, highlight the importance of integrating plural perspectives to capture the complexity of interactions with biocultural systems [54] [55]. For many communities, sentiments of relationality are of equal or greater importance than monetary value, and therefore, incorporating plural perspectives that account for the complexity of Indigenous Peoples' relationships to biocultural systems will be a key priority for any valuation of stocks, flows or benefits [5455].

Distributional layers on land management and ownership characteristics might help to reveal how natural wealth is distributed and how distributional effects of environmental policies might be anticipated using ecosystem accounting [56]. In ecosystem services accounts, considering distributional issues in the supply and use of ecosystem services and location of ecosystem assets could lead to more effective and economically efficient policy formulation [56]. It also permits the exploration of concerns relevant to environmental justice. This is particularly important for exploring connections between land inequality, ecosystem degradation, and wealth redistribution as a result of conservation or natural capital policies [56]. Therefore, distributional issues deserve to be more at the heart of ecosystem accounting, especially as this measurement tool becomes ever more present and integrated within the balance sheets of nations [56].

Additionally, pluralistic valuation approaches, emphasizing the societal and cultural dimensions of ecosystems over market-based metrics, offer pathways for identifying the best policy options without reducing nature to a mere commodity [57]. While monetary valuation methods are useful for comparative analyses, it is important to acknowledge differences in valuation methods and how they affect the comparability of values between different types of ecosystem services and between different scales and geographic contexts [58]. Finally, well-being, though briefly mentioned in the SEEA-EA framework, lacks consideration in terms of metrics within this accounting system, leaving a critical gap in connecting ecosystem accounting with holistic human welfare [30].

**14. Challenge: There is the need for an explicit consideration of intermediate ecosystem services, along with clear rules for allocating inter- and intra-ecosystem service flows and for avoiding double counting.**

SEEA states that ecosystem accounting should focus on final ecosystem services. However, to better understand the specific contributions of individual ecosystems to the economy and improve ecosystem management, it is important to also consider key intermediate services [59]. These intermediate services often represent flows that are part of an observable and material chain leading to final ecosystem services and associated benefits, thereby allowing a more complete recording of contributions by individual ecosystem assets and their ecological interactions [59]. This extended scope would enable the recording of dependencies, such as when a final ecosystem service supplied from one ecosystem asset relies on interactions with other ecosystems across different locations [59].

Delineating the chain of ecosystem service flows remains a significant challenge [60], necessitating an explicit consideration of intermediate steps and ecosystem services. Intermediate ecosystem services can manifest in different ways, such as "vertical" flows, where multiple ecological interactions converge into a single ecosystem service flow, or "horizontal" flows, where the same service is sequentially provided by different ecosystem types, highlighting inter-ecosystem flows [40]. Intra-ecosystem flows, occurring within the same ecosystem type, also require attention, as they contribute to maintaining ecosystem conditions and provide indirect support for human activities [40]. For instance, soil retention is an ecosystem service provided by most terrestrial ecosystems but is only accounted for as a final ecosystem service on cropland, where it is allocated to the agricultural sector [40]. In other cases, it remains an intra-ecosystem flow with diverse contributions [40]. To address these complexities, ecosystem service supply and use accounts should include entries that attribute part of the value generated by final services to intermediate services or record flows between ecosystems, thereby recognizing dependencies and avoiding double counting [59].

<u>Monetary Ecosystem Assets</u>

**15. Challenge: Modelling future ecosystem service flows entails interlinked difficult-to-make assumptions and inherits uncertainty of the data from other account tables, which is not reflected in monetary ecosystem asset accounts and their methodological framework yet.**

Modelling future ecosystem service flows faces significant challenges due to the need for detailed spatial-level data, while official projections are often available only at the national level [1]. Furthermore, there is ambiguity regarding whether specific scenarios should be modelled and, if so, which ones [1].

The common assumption that future prices for ecosystem services will remain constant is also problematic, as ecosystem capital scarcity could lead to upward pressure on such prices, potentially underestimating their value [61]. Similarly, it is doubtful that the future flow of income for each ecosystem service, physical flows and exchange values will remain constant over time [61].

Predicting future ecosystem service flows requires understanding the dynamic and non-linear relationships between asset condition and ecosystem service capacity, which are influenced by management practices, patterns of use, and changes in ecosystem condition, all of them difficult (or not always possible) to be determined [62]. This complexity is compounded by the potential impact of climate change, which could increase the value of some services while reducing others, further altering exchange values [63]. However, such changes are rarely integrated into monetary valuations [63]. Moreover, ecosystem dynamics often exhibit non-linear behaviors, including thresholds, multiple steady states, and feedback mechanisms, making the forecasting of service flows and the valuation of ecosystem assets highly uncertain, particularly under non-sustainable flow conditions [42].

The common net present value approach for calculating the overall value of ecosystem assets assumes no future degradation and consistent income flows, which is not a realistic assumption [64]. As a first issue, valuing future flows of ecosystem services involves predicting future conditions, quantifying uncertainties in line with attitudes toward risks, inferring preferences of future generations, and including non-use values, being all of this difficult to determine [32]. Additionally, as a second issue values related to ecosystems and biodiversity may emerge from socio-political processes rather than be pre-existing, requiring consideration of valuation in this context [32]. As a third issue, monetary valuation methods employ diverse concepts such as market price, opportunity cost, and willingness to pay, which are not always compatible. Some scholars suggest that the diversity of monetary valuation, instead of being additive, can provide a rich, inclusive information system for varied policy needs [32].

In urban contexts, few attempts have been made to estimate monetary values for ecosystem assets [14]. These efforts are hampered by high uncertainty surrounding future service flows, driven by climate change impacts, socio-economic trends, consumer preferences, and technological advancements [14].

**16. Challenge: Robust agreed-upon principles for estimating and allocating value loss due to ecosystem degradation are still missing in monetary ecosystem asset accounts and its methodological framework**

The allocation of ecosystem degradation poses significant challenges, primarily in determining whether to attribute the degradation to the economic and human activities that cause it (activity-based allocation) or to those incurring the costs of degradation (receiver-based allocation) [35]. For activity-based allocation, it is necessary to establish the relationship between the economic units responsible for degradation and the actual ecosystem service flows [35]. Degradation affects future flows of ecosystem services, but estimations often assume constant flows, even when there is degradation, overlooking the dynamic nature of ecosystem service supply [61].

A critical problem lies in estimating the value of degradation, as using the unpaid cost of restoration as a proxy is inconsistent with standard methods for valuing depreciation or fixed capital consumption [50]. Additional concerns include determining whether liabilities should be recognized and ensuring coherent accounting entries to avoid double counting, such as recognizing both asset value declines and liabilities for the same event Ogilvy et al., 2018). Furthermore, international accounting standards (IAS) and the System of National Accounts (SNA) consider it inappropriate to treat ecosystem degradation analogously to depreciation or capital consumption [50].

Some scholars indicate that the valuation of ecosystem degradation requires a sequenced approach, being the valuation of individual ecosystem services just one of the steps [62]. For estimating ecosystem degradation, a variety of assumptions must be made about the relationship between income (for ecosystem accounting the income refers to the ecosystem services) and capital (i.e. the ecosystem assets) [62]. However, it is only by working through these steps that a legitimate measure of degradation can be derived to be integrated with standard measures of depreciation and used to adjust aggregate income measures such as GDP [62].

Measuring degradation itself is complex, as it can involve changes in net present value (NPV) or the sustainable capacity of an ecosystem asset [62]. Sustainable capacity, reflecting the aggregate value of ecosystem services, diminishes as ecosystem conditions decline, offering a potential way of estimating degradation [62]. However, degradation is not merely the change in NPV, since the total change in NPV can be decomposed into changes due to human activity, changes due to price variability (revaluation), and changes due to natural processes and events (including degradation) [42].

For accounting purposes, degradation should be defined to allow its costs to be deducted from income earned through ecosystem use, aligning with depreciation concepts [42]. However, practical attribution to specific actors (e.g., landowners, industries, or governments) is challenging due to long-term impacts, cumulative degradation (ecological debt), and ecological thresholds [42]. Furthermore, allocating degradation to the appropriate economic units is complicated by spatial and temporal mismatches between degradation causes and effects, as well as the impacts on various ecosystem services, then, the attribution of overall impacts becomes quite complex [52].

The current SEEA-EA framework relies on simple causal representations linking pressures, condition changes, and impacts, but in complex systems characterized by non-linearity and uncertainty local causal chains cannot longer be assumed [32]. Ecosystem service pricing often assumes sustainability, ignoring potential depletion or degradation costs [62]. Similarly, restoration costs should not substitute as degradation values since they do not equate to lost future income potential [62]. In practice, assumptions of unchanged future ecosystem capacities for regulating services and production are applied, which are unrealistic particularly under overexploitation scenarios [65]. Ultimately, agreement on robust methodologies is needed to account for degradation's economic, spatial, and temporal complexities within urban ecosystem accounting.

**17. Challenge: There is still not enough evidence and consensus on the appropriate asset life value(s) to be used in monetary ecosystem asset accounts.**

There is still insufficient evidence and consensus on the appropriate asset life value(s) to be used in monetary ecosystem asset accounts. For example, in a report Defra/ONS suggests a 100-year asset life to reflect the longevity of renewable natural assets, but this timeframe lacks justification [5]. Assuming that ecosystems 'produce' services for a 100-year period is somewhat arbitrary, but common discount rates applied renders values beyond this period negligible in terms of Net Present Value [61,64]. In fact, ecosystem accounting examples from both the Netherlands and the UK rely on assumptions of no future asset degradation, constant ecosystem service flows, constant prices, and a 100-year ecosystem asset life, but do not justify those decisions [63].

Some scholars also made evident that asset values depend heavily on management practices and assumptions about asset life. For instance, individual tree asset values in Oslo were calculated by discounting future benefit flows based on species-specific lifetimes under urban conditions [63]. However, this approach underestimates the value of trees that have not reached their climax growth phase and neglects the potential value of locations where replacement planting could occur after trees reach the end of their lifespan [63].

Different than a constant value, the SEEA EA framework recommends basing asset life estimates on recent patterns of ecosystem use, rather than on generalized assumptions about future sustainability or intended management practices [63]. However, in urban contexts, such as in the municipality of Oslo, recent municipal plans emphasize tree conservation and planting, suggesting that past management may not be a reliable predictor of future asset conditions or lifespans [63]. This will also occur in many other urban contexts in the European Union, especially after the target on increase of urban tree cover included in the Nature Restoration Regulation, making also the use of recent pattern of ecosystem use to establish the lifetime of ecosystem assets problematic.

**18. Challenge: There is still not enough evidence and consensus on the most appropriate discount rate(s) to be applied to future ES flows.**

Current approaches vary significantly between countries and contexts. For example, in the past, in the UK, ecosystem service values relied on Green Book guidance for project appraisal, aligned with the DEFRA/ONS approach, applying a graduated discount rate starting at 3.5% for the first 30 years, 3.0% for years 31–75, and 2.5% for years 76–100 [5]. In contrast, the Netherlands applies a rate of 2–3% depending on the ecosystem service type, with recommendations to use 3% for provisioning services such as agriculture and timber, and a lower rate for irreplaceable services [61 63 64]. Similarly, the Netherlands Environmental Assessment Agency (PBL) suggests considering increases in relative prices due to scarcity and substitution limits, resulting in an effective rate as low as 2% for some services, although the rationale for these decisions remains partly subjective [61] Schenau et al., 2022).

The broader debate on discount rates is shaped by varying recommendations, such as Stern's (2007) proposed 1.4% rate for climate change mitigation investments [63]. Meanwhile, the System of National Accounts (SNA) advocates using market discount rates for net present value (NPV) calculations, yet it remains unclear if these are suitable for non-market ecosystem services in accounting frameworks [42]. Some argue for aligning discount rates with market transactions, but non-market services, such as air filtration, lack equivalent market benchmarks. Consequently, applying public discount rates to non-market services might be appropriate, pending further academic and statistical discourse [42]. The challenge thus persists: achieving a consensus on determining discount rates that balance theoretical consistency with practical application across diverse ES contexts, in part because of a lack of evidence.

**Cross-cutting**

**19. Challenge: Varied and unclear policy uses of urban ecosystem accounts across multiple spatial levels (national, regional, local) create diverse requirements, risking the loss of coherent framing.**

Ecosystem accounts are still in the early stages of development and often lack a unified framework, which hinders their integration into decision-making processes [66]. A challenge for municipalities in adopting SEEA EA at local level is that policy priorities, their geographical scale and resolution are different from national accounts [35]. In fact, there is an inherent disconnection between the ecosystem accounting needs of local governments and those at the national level, as the policy issues differ across scales, and even regionally within local contexts [1].

Moreover, while some progress has been made towards urban ecosystem accounting -such as the inclusion of vegetation structure and information on public access to green areas- there remains a need for additional data to assess urban ecosystem services as a basis to inform land use planning and policy [8]. However, urban ecosystem studies often lack standardization, with methodologies, concepts, and tools being developed in isolation for different sectoral themes [67]. This lack of coherence prevents the comparability of results across sectors, and the potential for integrating ecosystem services into broader policy remains limited [67]. Standardization of urban ecosystem accounts would solve this, but it will require considering the multiple sectorial policy needs of urban contexts in a comprehensive manner.

The potential uses of ecosystem accounts are numerous but still largely speculative or anecdotal [68] [32]. To move toward institutionalization, ecosystem accounting requires a "policy pull"—a strong and consistent demand for information from users, which is currently lacking in most parts of the world [69]. This results in a gap between the production of ecosystem accounting data and its actual use in decision-making [69]. In fact, the development of ecosystem accounts for a while mostly focused on overcoming technical challenges without considering enough policy-driven demand [70]. For further actual use of accounts, more discussion and investigation are needed to identify and specify potential uses, validate them, and discuss their relative relevance for supporting the transition to sustainable societies [32]. As a consequence, ecosystem accounting is not yet widely used by local city planning institutions, although standardized accounts could help track trends in the provision of urban ecosystem services and inform climate-action plans [10].

To clarify the intended uses of ecosystem accounts are critical, as they influence their level of detail and development. This clarification requires reinforcing a co-construction framework complementing the "technical-push" orientation that largely dominated the development of accounts [32]. Many important discussions could not be solved rigorously without an explicit and specific identification of the main intended uses of the accounts and their context [32]. It would be different if ecosystem accounts were used for specific decision making at local levels rather than to provide broad indications about ecosystem (and corresponding economic) trends [71]. Moreover, the audiences for ecosystem accounts are still not clearly defined. It will be necessary to avoid the expectation that the accounting system alone will be able to meet the broad range of needs of all potential end users [71].

While there are some examples of ecosystem accounts supporting decision-making these are still limited and often not linked to a broader policy context [72]. Ecosystem accounting could be useful in the phases of policy design (policy formulation and decision-making), policy implementation, and policy evaluation in the policy cycle [72]. To improve their usefulness, there must be clearer communication about their intended policy uses [66]. Effective implementation of ecosystem accounts requires the mobilization of knowledge, integration of data across sectors, and assessment of ecosystem services [66].

Finally, in cases where urban ecosystem accounting pilots are being developed, such as in Australia, single urban ecosystem accounts are being developed in isolation from the others, without a consistent or unified approach to quantifying and valuing urban ecosystem assets [14]. As a consequence, this leads to no consistent or unified approach to defining, classifying, quantifying, and valuing urban ecosystem assets for the purpose of developing environmental-economic accounts [14]. There is a need for better coordination to develop a system that supports both top-down disaggregation from the national scale and bottom-up aggregation from local scales, ensuring that ecosystem accounting can serve multiple beneficiaries at various policy levels [73].

**20. Challenge: There are still missing complete and consistent examples of ecosystem accounts that can serve as practical guidance.**

Many pilot experiences fail to meet the criteria of condition accounts as outlined by SEEA EEA or SEEA EA rules [74]. Besides this issue, there were only three urban ecosystem accounts identified to date [74]. In addition, the development of accounts for a broader range of ecosystem services has been indicated as essential, as each service exhibits unique characteristics that require tailored modeling and accounting approaches [75]. Furthermore, it is also indicated that additional studies on the economic valuation of ecosystem services, such as monetary ecosystem services accounts or monetary ecosystem assets accounts, could highlight the importance of ecosystem accounting in informing natural resource management decisions [58]. The last point could not only increase in the first place the number of examples of specific ecosystem accounting tables, but also promote the generation of more examples that would help to achieve consistent ecosystem accounting examples serving as robust practical guidance.

**21. Challenge: Ecosystem accounts still lack an explicit assessment and reporting of uncertainties, along with an agreed-upon methodology.**

Guidelines for ecosystem accounting remain insufficient, making it difficult to address uncertainties in the models and valuation processes [1]. Forecasting future flows of benefits and market prices/values, for example, often relies on assumptions such as constant ecosystem service production and fixed prices over long periods (e.g., 100 years), which are inherently uncertain [5]. Additionally, monetary valuation models are influenced by the lack of data availability and uncertainties in biophysical models, with variations in model accuracy depending on the ecosystem service being considered [61 61].

In some cases, estimated ecosystem service (ES) indicators and aggregate monetary asset values lack sufficient accuracy and reliability, which limits their utility for detecting trends in ecosystem asset values (Cimburova & Barton, 2020). In urban ecosystem accounting, it has been illustrated as problematic when modeling assumptions yield values that surpass the observable changes in urban canopy cover during accounting time steps [76]. Furthermore, the data used for accounting purposes is often sourced from government and research agencies, but the accuracy of these datasets may be questionable, further complicating the estimation of ecosystem service supply and asset value [61]. The complexity of ecosystem dynamics, including ecological thresholds and stochastic events, presents another issue for accounting uncertainty in ecosystem accounts [61]. Although these dynamics could theoretically be integrated into ecosystem accounts, the uncertainty surrounding their occurrence and impacts makes it difficult to include them accurately [61].

Moreover, the models and ecosystem services maps used for accounting are not well-verified for accuracy, particularly at the local or municipal level [61]. Differences in input data, modeling methods, and the level of spatial explicitness all contribute to variability in accuracy. Studies often rely on secondary land use or land cover data and value transfer approaches, which may lead to substantial errors, especially in urban ecosystem accounts where landscapes are heterogeneous [61 23]. The lack of validated models in ecosystem services assessments further exacerbates this issue, as combining different types of data (with different degrees of spatial explicitness and spatial variation) can introduce errors that are difficult to quantify [51].

As a consequence, the persistence of uncertainties (non-assessed and reported) in both physical and monetary estimates of ecosystem services hinder the integration of ecosystem accounts into broader policy areas [63]. This is compounded by changes in ecosystem conditions. For example, in the case of air pollution levels, expected changes in conditions are not accounted for when estimating future ecosystem service flows like air purification, potentially leading to overestimated asset values if pollution levels are assumed constant, instead of being under decreasing trends due to technological changes and new air quality policies such as in the case of Europe [63]. Ecosystem accounting therefore requires explicit attention to uncertainty and sensitivity analysis, including the publication of uncertainties and the selection of the most appropriate modeling approaches [42].

There is also an ongoing need for discussions regarding the acceptable level of transfer error in ecosystem accounting, particularly when value transfer methods are applied [77]. These methods often have error rates comparable to those found in standard national accounting, but more information is needed to improve the accuracy of transfer, validity, and adjustments for error minimization [77]. Moreover, ecosystem extent and condition accounts must better incorporate quantitative measures of error or uncertainty, as measurement errors and systematic errors in model representations are significant challenges [78]. In the case of extent, the misclassification of ecosystem extent due to dataset inconsistencies further complicates the reporting of errors or uncertainty [3]. In fact, sometimes recorded additions and reductions in extent accounts are not real, but an artifact from classification errors in the ecosystem extent maps [35]. Finally, the quantification and valuation of regulating services, such as those provided by coastal and marine ecosystems, suffers from high scientific uncertainty, making reliable accounting even more difficult [73].

**22. Challenge: Expertise, time and resource demand generate practical barriers for the implementation of urban ecosystem accounts.**

It has been indicated by some experts that developing the first set of accounts is a knowledge- and resource-intensive task, requiring at least 15–20 person months over 2–3 years, with additional funding allocated for primary data collection (United Nations, 2020). This process necessitates an independent technical group to address the highly technical and rapidly evolving nature of the work until the accounts are streamlined and automated (United Nations, 2020).

Despite advancements in GIS tools that bring ecological modeling into statistical offices, professionals must customize and adjust variables, ensure outcomes are meaningful, and interpret results appropriately, which further complicates the process (United Nations, 2020). Moreover, ecosystem accounting models are complex, requiring specialized expertise that is not easily accessible to practitioners or policymakers [75]. Compiling a full suite of ecosystem accounts demands substantial data and multiple biophysical models, which vary in scope and spatial detail depending on budget, technical capacity, and data availability [35].

While the SEEA EA provides a statistical framework, further methodological guidance is still needed; organizations such as UNSD and Eurostat are developing guidelines and technical reports to address biophysical modeling and ecosystem service valuation [47].

Among specific countries, Nordic countries illustrate the resource-intensiveness of this endeavor, highlighting national coordination challenges due to diverse input data formats, the spatial nature of much of the data, and the lack of GIS experience among statisticians and economists [58]. This underscores the need for skill development and the employment of personnel with other types of skill sets different than those of traditional employees of statistical agencies [58]. At the municipal level, local personnel and resource constraints, coupled with varying priorities among municipalities, further hinder the uptake of ecosystem accounting compared to the national level [35]. Establishing mapping and accounting workflows and applications requires significant initial investment. Therefore, to facilitate regular implementation at the municipal level, regional or national agencies need to develop common approaches [35].

**23. Challenge: There are still gaps, and lack of consensus, regarding suitable input data, data quality frameworks, and generalizable models for use in ecosystem accounts.**

In part this issue stems from the limited availability and inconsistencies of data across different spatial and temporal scales. Statistical data, often collected and reported by administrative units, must be spatially disaggregated to provide grid-based results needed for ecosystem accounting (United Nations, 2020). However, this disaggregation process may not always be applicable, and account compilers must decide when such an approach is more effective than direct modeling (United Nations, 2020). Furthermore, the datasets commonly used for ecosystem accounting often lack temporal, spatial, and unit coherence, making it necessary to build capacity in statistical reporting and data collection at all levels [60]. The scarcity of accurate data frequently leads to the use of rough estimates, underscoring the need for transparent data management policies [60]. Another challenge is that the data needed for ecosystem accounts are typically not collected by statistical offices, but rather by many different ministries and agencies for monitoring purposes [79]. As a result, these datasets are often irregular, inconsistent with standard statistical classifications [79].

In addition to issues related to data availability, the resolution of models can significantly influence ecosystem service (ES) valuations. Sensitivity analyses have demonstrated that the spatial resolution of models can impact the valuation of ES, indicating that further attention is required in this area [80]. Furthermore, data availability often does not match the required time frames or spatial resolution, complicating the development of consistent accounts for a representative time series [75]. This challenge is especially pronounced in urban ecosystems, where modeling ecosystem services is complex because of the heterogeneous nature and the variety of sizes, urban morphologies, climates, and socioeconomic characteristics of various beneficiaries in different cities [10]. Factors such as scale, accessibility, heterogeneity within ecosystems, and budget limitations all affect modeling feasibility [46]. Additionally, urban ecosystem services at national scales face challenges due to the lack of comprehensive data [10]. Moreover, relationships underlying ecosystem service provision may vary by city, depending on its size, density, or climate zone [10]. This complexity is compounded by data limitations that hinder the ability to address management questions, particularly regarding ES trade-offs [81].

Another critical issue is the lack of consensus on the appropriate methods for valuing ecosystem services, particularly for services like free access recreation [44]. There is also a notable gap in the integration of environmental and economic data, particularly the lack of spatially referenced economic data, which limits the effectiveness of ecosystem accounting [72]. Standardization of methodologies across countries would improve the suitability of data for regular updates of ecosystem accounts [82]. Yet, the current methods for estimating ecosystem services often contain biases due to assumptions regarding service use or supply [83], and the underlying valuation methods are not always suitable for services lacking market prices, further complicating the process [58]. There is also a need for standardized design and reporting in primary valuation studies to facilitate value transfer and accounting purposes [77]. This includes clear identification of components in the Supply and Use Tables for ecosystem services, which would require a reporting protocol for primary studies to ensure reliable input data [77]. Important information would include sensitivity (i.e., parameterization where possible) of values to ecosystem extent, condition and relevant spatial variables, population characteristics and institutional contexts, as well as standardization in units of measurement [77].

In terms of modeling, many generalized models, such as ARIES and InVEST, face challenges when applied to large heterogeneous environments, despite their computational feasibility. These models' parameterization and calibration across multiple contexts remain difficult [84]. Maintaining internal coherency and comparability is one of the main challenges of ecosystem accounting. In fact, accurate ES accounting is still very challenging and demanding in terms of data: substantial work is needed to adapt and test methods that can be and remain consistent with national accounts [85]. Moreover, ecosystem accounting often requires complex models with high data and computing needs, which may not always be available or feasible. The lack of consistent and reliable data complicates the development of ecosystem accounts, particularly for cases such as ecosystem condition, ecosystem capacity for resource provision [65]. For instance, past pilot ecosystem accounts at the Italian level, faced challenges mainly related to data availability and time misalignments, further hampering the creation of consistent time series [86].

Lastly, accurate ecosystem accounting requires efforts to address data shortcomings, such as mismatched time periods and spatial inconsistencies, which hinder the development of reliable time series [35]. To overcome these challenges, closer collaboration between scientists from diverse disciplines and decision-makers is needed, as ecosystem accounting relies on the expertise of statisticians, land managers, economists, and ecologists, among others [51]. Furthermore, research should prioritize the analysis of models with respect to their application requirements and potential, considering factors like spatial scale, input data requirements, and computation capacity, to enhance their applicability in ecosystem accounting [87]. Additionally, policy agencies often lack familiarity with how to use the information generated from ecosystem accounts, which creates barriers to their effective use in decision-making processes [79]. Thus, fostering capacity building across all stakeholders is key to overcoming the data and modeling challenges in ecosystem accounting.

**24. Challenge: There is still a need to establish common principles and practices for developing data infrastructures for ecosystem accounting, ensuring interoperability and data sharing among stakeholders.**

Regular data updates, including digital spatial data, require promoting inter-institutional coordination and collaboration through both top-down and bottom-up initiatives, alongside the development of national spatial data infrastructure for ecosystem accounting [60]. Institutional challenges often arise when integrating data from different agencies, as datasets may exist in incompatible formats, or there may be reluctance to share them [35]. Enhanced collaboration between government institutes holding disparate datasets could improve data integration and foster greater commonality in terminology and definitions [35].

Centralized, replicable data and model management strategies, incorporating code repositories and interoperability-focused approaches, are essential to ensure the reliability and replicability of account compilation [69]. This is particularly crucial for long-term ecosystem accounting, where accounts will need to be recompiled as data or modeling advances [69]. In supranational accounting under a legal framework, data harmonization is vital to enable the harmonized collection of comparable data [69]. Furthermore, ecosystem accounting studies are often partially incomplete, with limited transparency in methods for compiling extent and condition accounts [78]. Enhancing data accessibility and improving the analytical methods used for these accounts would enable accurate comparisons between accounts and increase the likelihood of achieving data interoperability and reusability [78].
