## Supplementary Information 4 for "Advancing the global statistical standard for urban ecosystem accounts"

**Supplementary Information 4: Description of lessons and approaches**

This supplementary information provides a detailed description of the lessons and approaches based on the information collected during the systematic and thematic review. Lessons point either to univocal solutions or to key constraints when addressing specific challenges, while approaches represent potential alternatives that occasionally appear contradictory in the literature.

The lessons and approaches are organized by challenges, as presented in Table 2 of the manuscript. Deliberately, the authors have limited their own interpretation in describing the lessons and approaches below, ensuring that they remain differentiated from the discussion and proposals presented in the main manuscript. However, in some instances, brief notes or clarifications have been included where necessary. While these additions inevitably reflect the authors' interpretation, they aim to enhance clarity and facilitate comprehension of specific lessons and approaches.

**Ecosystem Extent**

1. **Challenge: There is still no univocal distinction between urban ecosystem assets and artificial surfaces.**

**Approach: A dominant approach in Europe considers that urban ecosystem assets are built up land where people live and its surroundings, including semi-natural and natural patches.**

Urban spaces feature high population density and a concentration of human activities and artificial assets [1 2]. They can be depicted only as the densely built-up land where people live (settlements and other artificial surfaces) [2]. In a broader sense, they can be defined as both the densely built artificial zones and their surroundings, which also include natural and seminatural components [1]. This second perspective corresponds to urban ecosystems, where built-up areas are one of the components of the ecosystem [2].

For accounting, to distinguish between urban ecosystem assets and artificial surfaces, two complementary approaches can be proposed: a general approach and a thematic approach [2]. The first focuses only on highly artificial land and omitting overlaps with other ecosystem types (settlements and other artificial surfaces) but does not fully represent the overall urban ecosystems [2]. The second focuses on urban ecosystems defined as cities and their surrounding socio-ecological systems, which includes a mosaic of anthropogenic, natural, and semi-natural land covers [2].

### 2. Challenge: There is still no univocal procedure for the spatial delineation of urban ecosystems and their accounting areas.

**Approach: There are already several alternatives for delineating urban ecosystems.**

Various methodologies have been proposed for the spatial delineation of urban ecosystems and their accounting areas:

- Built-up areas plus buffer: the urban boundary is defined using a variable-sized buffer around each polygon representing built-up areas, incorporating adjacent non-built-up patches such as large parks based on the size of the polygon [3].
- The Degree of Urbanisation: it offers a classification along an urban-rural continuum based on population size and density thresholds [4]. It captures three mutually exclusive classes (cities, towns, and rural areas), being a classification that can be applied globally, permitting comparisons.
- Functional Urban Areas: It can complement the Degree of Urbanisation, delineating urban ecosystems as cities (as defined in the degree of urbanization) plus their surrounding commuting zone [4] [5].
- Administrative boundaries: They can be useful for defining urban ecosystems, using standard census geographies and merging adjacent municipalities based on population and economic integration thresholds [1] (Taubenböck et al., 2022). This delineation of urban ecosystems is significant for jurisdictional purposes and with local data compilation practices [5] (Taubenböck et al., 2022). However, it often fails to reflect the actual physical extent of urban areas, especially in large cities that extend beyond these limits [7].
- The morphological approach: It delineates urban ecosystems based on continuous built-up land cover classes and population thresholds [8] [9]. There are two approaches in Europe, Morphological Urban Areas (650 inhabitants per km2 and with a minimum population exceeding 20,000 inhabitants) and Urban Morphological Zones (continuously built-up areas derived from Corine Land Cover with a minimum population of 10,000 inhabitants) [8] [9]. The morphological approach is particularly useful for studying land-use patterns and infrastructure planning [8].
- The ring-based approach: it delineates urban ecosystems by making use of consecutive rings of 1km width from the city center outwards [10]. The outer ring limiting the urban ecosystem should: i) be large enough to incorporate continuous urban areas, ii) not include isolated small cities far from the main urban area [10].
- The grid-based approach: it offers a more localized and porous mapping, utilizing a consistent spatial grid to determine urban versus rural areas based on building density, population density, and share of certain building types (Taubenböck et al., 2022).
- The urban gradient approach: It defines urban ecosystems through analysis of anthropogenic gradients, generated as a consequence of the creation of human settlements [11]. Urban gradients can be defined through broad and specific measures of urbanization [11]. The first can be easily measured (e.g., density of people, density of dwellings, shared of built surfaces) but

might capture a complex range of conditions. The second are more difficult to measure but might provide a more precise measure of urban characteristics and their likely direct influence on ecological responses [11].

**Lesson: The delineation of urban ecosystems influences consideration of transboundary effects (impacts and dependencies) in ecosystem accounts.**

The delineation of urban ecosystem boundaries for accounting purposes is usually framed only thinking in assessing the contribution of assets located in urban areas and the derived services and benefits consumed by population living there [5]. However, part of the population that consumes urban services and benefits might live outside urban areas and simultaneously "non-urban" assets might contribute to the supply of services in urban ecosystems [5]. Then, to better consider impacts and dependencies (i.e., functional relationships) of urban ecosystems, the delineation of its boundaries for accounting purposes might consider an extension of its limits [5]. It might be also necessary to determine the boundaries per ecosystem service [5] .. As an alternative to the adjustment of boundaries, it could be included an assessment of the dependency of urban ecosystem assets on the status and productivity of non-urban assets located near or far away [5]. Accounting for transboundary effects could provide a more comprehensive and accurate urban ecosystem accounting framework by better highlighting dependencies and impacts.

**Lesson: Aspects influencing the delineation of urban ecosystem boundaries might not be entirely objective.**

Regarding delineation of urban ecosystems, research shows that there are often no clear, incontrovertible facts, but only different ways of conceptualizing, measuring and interpreting the urban ecosystems (Taubenböck et al., 2022). The definition of urban ecosystems usually depends on its conceptualization, the data, the spatial units of measurement, the scale of the accounting (local, regional national) and the variables or the thresholds applied (Taubenböck et al., 2022) [5]. It also depends on the intended use of the urban ecosystem accounts [5]. For example, if the intended use is informing specific urban policies or plans, the entities of reference in which those are applied become relevant when defining limits of urban ecosystems [2]. This means that the definition of accounting limits, but also specific reporting units, for urban ecosystem accounts are crucial elements influencing their intended use, and they may vary depending on the policy focus. [2]. In synthesis, research seems to support the idea that there is no best concept, set of data or methods, but different approaches (Taubenböck et al., 2022). As a result, it has been also suggested that it might be better to combine multiple approaches when delineating urban ecosystems, providing a probability-based range of the degree of an area of being an urban ecosystem (Taubenböck et al., 2022).

3. **Challenge: At a global level, urban ecosystems still lack an ecologically sound, agreed-upon classification of homogeneous ecosystem areas, along with the principles to develop it.**

**Approach: There are already several alternatives for dividing urban ecosystems in homogeneous groups.**

Classifying urban ecosystems into clusters of homogeneous units should help in the definition of meaningful reference levels for ecosystem condition variables within each urban ecosystem [2]. Various ways of dividing urban ecosystems in groups have already been proposed:

- Division by biogeographical units. This is considered frequently in the case of natural ecosystems, but it might be unsuitable for anthropogenic ecosystems such as agroecosystems and urban ecosystems [2]. Instead, it might be more suitable considered together with land use, management and socio-economic factors [2].
- Division by urban ecoregions. These are regional clusters that share some similarities with the use of biogeographical units, but which are defined not only based on macrofactors such as climate or vegetation, but also on urban topology [12]. It may facilitate a ecologically robust upscaling from individual cities to regional and global levels. [12].
- Division by urban growth patterns. It could be based on clusters of intraurban horizontal and vertical growth patterns (e.g., stabilized, outward, mature upward, budding outward, and upward and outward) [7 13]. It could be based only on horizontal growth patterns (e.g. Expansion, sprawl, polycentrism and densification/coalescence) [14]. Previous studies showed that cities within a geographic region usually share a similar urban growth cluster [13].
- Division by morphologic-spatial composition and configuration. Previously most experiences have based this characterization on the following dimensions: compactness, centrality, form complexity, porosity, density and other configuration characteristics [15 16]. Some of those experiences have considered only some dimensions, such as compactness [17] or compactness and centrality [10]. There are alternatives that bases this kind of urban characterization in the following factors: i) the average size of the cities; ii) the share of built vs non-built land cover or local climate zone classes (assessing intensity of human use); iii) the distribution of land cover or local climate zone classes within the urban areas (assessing spatial configuration); iv) building volume density gradients (assessing distribution of the usage of space) [9]. Finally, in some cases researchers have proved cross-sectional correlations between urban form and economic [7 15]. This might mean that characterizing cities by urban form could inform on underlying socio-economic factors and dynamics.
- Division by clustering of mixes of land cover types. In SEEA EEA, land cover has been frequently used as a proxy for ecosystem types [18]. Then, grouping by grouping of land cover mixing or dominances might be also considered for splitting urban ecosystems in groups.
- Division by connectedness with other urban ecosystems. The social, economic and ecological impact/relevance of an urban ecosystem does not depend only on that system itself, but also on its relationships with other urban ecosystems [19]. To characterize the importance of an urban ecosystem (as a node) in terms of its connectedness with other urban ecosystems (other nodes), the following metrics are commonly used: degree centrality, betweenness centrality, closeness centrality, eigenvector centrality, local clustering coefficient, local dominance. This approach can be valuable for clustering purposes [19].

Besides division by just one of the above approaches, some researchers also point to consideration of several of them (e.g., climatic, urban size and configuration, land cover, and socio-economic characteristics) to ensure homogeneity from several perspectives [2] EW(Schwarz, 2010).

**Lesson: Some common metrics are used when grouping urban ecosystems based on morphology and spatial composition and configuration.**

For characterization and grouping of urban ecosystems based on morphology and spatial composition and configuration there is a list of common metrics used for specific dimensions:

- Complexity (irregularity of patch shapes): landscape metrics such as area weighted mean shape index, edge density and area weighted mean patch fractal dimension is common [7 15].
- Centrality: usually measured using the distance of dispersed patches to the central patch [15].
- Compactness: landscape metrics such as compactness index, largest patch, class area, compactness index of the largest patch, number of patches referring all to patches classified as urbanized land or built-up area [7 15 17]. Those metrics are interested in aspects such as size and the level of clumping of individual patches of urbanized land.
- Porosity: measured through the ratio of open space compared to the total urban area [15 16].
- Density: measured through population density, which sometimes could be generic (e.g., with respect the entire urban ecosystem) or specific (e.g., with respect to the built-up areas inside the urban ecosystem) [15]. It have been also characterized with measures of employment density [17]. In both cases, it could inform on the intensity of urban land uses by humans [17].
- Size. It is commonly measured in relation to size of built-up area or size of human population.
- Composition: commonly measured as the proportion of built-up land cover classes with respect to the rest [15].
- Configuration: diversity indexes such as Simpson's index applied to land cover classes are commonly used [16]. Also, metrics such as core area of built-up areas or adjacency of built-up areas to other land cover classes [16].

There are other metrics that are less used such as topography, GDP, specific economic characteristics, historic legacy (e.g. Colonial influence) that might also influence morphology, spatial composition or configuration of urban ecosystems, being relevant to characterize them [9].

In general, studies use landscape metrics (or similar geospatial metrics that might go beyond classic landscape metrics), spatial statistics (e.g. Regression metrics, spatial autocorrelation, evenness of land type distribution), and some socio-economic variables [14].

Despite the common use of the metrics and the dimensions introduced above, in urban science it is not yet agreed which dimensions and metrics/features (including landscape metrics) are most suitable for the characterization of urban ecosystems [17]. In fact, some criticized that the use of metrics until now has been quite arbitrary, together with the selection of the scale of observation, the spatial reference systems used and the geographical boundaries [17]. In fact, some studies compare urban ecosystems among each other without a previous clear theoretical basis. Then, results are constrained to the cases compared, being their transferability and explanatory power limited [17].

### 4. <u>Challenge: Urban ecosystems still lack a common ecologically sound classification of the fine-scale patches (assets) that compose them.</u>

**Approach: There are already several alternatives for classifying the fine-scale assets that compose urban ecosystems.**

The ecology of cities paradigm suggests disaggregating urban areas into fine-scale assets (patches) of both built and non-built components, which are essential for urban system functioning [21]. Various methodologies have been proposed for the classification of fine-scale assets in urban ecosystems:

- Land use/land cover classes. This is a common approach, especially under the ecology in cities paradigm and which is commonly used in urban planning. Making use of land cover classes could be used to represent urban sub-systems of homogeneous character [2]. However, it tends to simplify developed areas into a homogeneously built matrix, failing to capture the fine-scale spatial heterogeneity [22] [21]. Additionally, built and non-built components tend to be split in different classes, when their mixing or integration is a fundamental characteristic of urban ecosystems [22] [21]. Moreover, some researchers indicate that land-use and land-cover classifications often mix cover and use, but differentiating both is relevant to test the link between heterogeneity and ecological functioning [22]. Maintaining the differentiation permits to use land cover as an independent variable for explaining use but also ecological processes/functions (e.g., nitrate export, carbon sequestration) [22].
- Gradient of urbanization. This approach shows internal assets of urban ecosystems classified in zones according to a gradient of high urbanization to low urbanization [23]. Urbanization gradients are essentially anthropogenic gradients, in which the major environmental changes are: (1) creation of new land-cover; (2) alterations to the chemical and physical environment; (3) the creation of new assemblages of organisms and (4) alterations to disturbance [11]. For this classification demographic and physical measures of urbanization are commonly used [11]. Among the variables to represent urban gradients the following are common: population density index, road density index, urban forest cover (including all green/vegetated structures), sealed surfaces and socio-economic status of the population [23]. Additionally, land shape metrics have also been proved relevant in some cases [11]. The above variables are usually considered as broad measures of urban gradient, but specific measures can also be used [11]. Among examples using urban gradients, the Atlas of Urban Expansion making use of this approach offers a classification of fine-scale assets in five categories: urban (50% or more urban pixels inside a 1 km2 buffer), suburban (25–50% urban pixels inside a 1 km2 buffer), rural (less than 25% urban pixels inside a 1 km2 buffer), captured open spaces (non-urban pixels like vegetation, fully surrounded by urban and suburban pixels), rural open space (those that are not fringe or captured), and water [7]. The Urban Atlas expansion is an example of making use of land use or land cover classes to define urban gradients. In fact, Land-use or land-cover are among the most commonly used measures for developing urban gradients [11].
- Multi-level classification. Among the approaches used a few splits the fine scale asset using multiple levels of division along a scale of detail [24]. First, using classifications of land based on independent attributes such as spectral reflectance, land use, and population density [24]. Second, interrelational attributes capture the spatial relationships, configurations, and arrangements between multiple elements, useful to explain flows of matter energy and information [24]. Third, splitting pieces of the urban ecosystem in stands (like forest stands) of structurally distinct sub-regions formed by combination of shared independent and inter-relational attributes [24]. Finally, splitting larger areas in stand mosaics (like landscape mosaics), which describe the composition and configuration of specific stands inside a larger region [24].
- Built-up area density. A few approaches assumed buildings and their density are the key factor, sometimes considered together with population density [10] [25]. For example, some classifications split urban ecosystems internally based on characteristics such as building and population density, building's year of construction, settlement's vertical profile, building's construction material [25]. Others just split the urban ecosystems into large sub-regions based on the density of the built-up area. For example, splitting urban ecosystems into four classes,

urban core, inner urban area, suburban area, and urban fringe according to 75%, 50% and 25% urban built-up area densities [10].

- Cells of mixed land cover + building height (STURLA). This approach splits fine-scale patches in regular cells making use of a grid, in which class is defined based on the share of tree canopy, low plants, soil, water, paved surfaces and building height types (low rise [1-3 stories, midrise [4-9 stories] and high rise [>9 stories]) [26]. The premise of this approach is that the relationship between urban structure and ecological functioning requires an examination of the complexity of the urban structure, in particular how urban internal assets are organized [27]. Since the third dimension might be highly variable in urban ecosystems and could influence other abiotic and biotic factors, it is assumed necessary to consider it explicitly in the classification. Applications of STURLA have shown that landscape structure classes are related with land surface temperature and phylogenetic diversity of the atmospheric microbiome [28]. Researchers advocating for STURLA indicate that some processes such as cooling effects derived from building shadowing may be missed in classification systems that are not compositional and explicitly integrate building height [28]. Then, these other systems might miss interactions that may influence ecological and environmental parameters, such as LST [28]. In fact, as a novelty, STURLA offers an automated composite functional classification of urban structure that includes the vertical dimension, and thus can be applied to wide geographic regions systematically [29] [27] [30] [31]. Moreover, STURLA works at small spatial scales (120 m2), which are still meaningful for urban planners making pixels geographically meaningful for neighborhood development projects [28]. However, since STURLA makes use of a regular grid of cells, if grid size, location or orientation were shifted it may change the relative proportions of within class elements [27]. Moreover, groups have a range of variations between components, then cells belonging to the same class are subject to within class variation [27].
- Patches of mixed land cover + building height (HERCULES). The HERCULES approach splits urban ecosystem patches based on six landscape features: coarse-textured vegetation (trees and shrubs), fine-textured vegetation (herbs and grass), bare soil, pavement, buildings, and building typology [32]. Water is represented by the absence of the other elements [32]. The proportional cover is divided into four ranges: (0) absent, (1) present to 10% cover, (2) 11–35% cover, (3) 36–75% cover, and (4) > 75% cover [22]. Building typology, has five recognized types: single structures in rows or clusters, connected structures (share a wall or are associated with multiple walkways while sharing the same roofline) mixed, (with multiple wings, connection by courtyards or arcades, or a group of buildings with different structural footprints), high rises (between 4–10 stories), and towers (greater than 10 stories) [22]. To accurately reflect ecosystem function, HERCULES requires patches to be a minimum of 20 m in two orthogonal directions, preventing small or isolated features such as individual city lots from being classified independently [22]. In HERCULES, the type of vegetation, surface material, and buildings are hypothesized to influence ecosystem function because of their differential influence on the amount and distribution of organisms, materials, and energy. Similar to STURLA, it disaggregates urban areas into specific hybrid patches that include both built and non-built components, whose integrated consideration is crucial for studying urban system's functionality [21]. It also distinguishes between structure, use and function, making the classification of fine-scale assets an independent variable for studying ecological functioning [22]. In fact, the structural features of hybrid patches can reflect biological processes, such as plant invasion, survival, and succession, but also can express social criteria, such as zoning and

construction regulations [21]. In synthesis, the key characteristics of HERCULES are that it: (1) integrates human and natural components of the landscape; (2) recognizes that features can vary independently of each other; (3) accounts for all combinations of elements in the landscape; (4) has greater resolution in distinguishing categories; and (5) distinguishes between structure and function [22].

- Local climate zones (LCZ). This approach represents a standardized and harmonized framework for classifying urban ecosystems based on universal, measurable parameters of urban form. The system includes ten built types (lczs 1–10) and seven land cover types (lczs A–G), each with specific climate-relevant physical parameters (Huang et al., 2023). These 17 distinct patterns are composed of buildings, roads, plants, soils, rock, and water, each arranged uniformly and distinguishable on city maps or aerial photographs [34]. Lczs categorize urban landscapes using parameters such as density, building size, and building height for built areas, and thematic elements like trees or open spaces for non-built areas [9]. To implement lczs, two types of basic spatial units are used: parcel units (e.g., lot area polygons, urban blocks) and grid units, which divide the city into square cells. Parcel units typically have a minimum radius of 200–500 m, while grid units' appropriate size can be determined empirically, through testing, or based on building height spatial autocorrelation (Huang et al., 2023). LCZ was originally developed for urban climate studies, specifically to address the confusion in differentiating urban and rural areas in Urban Heat Island (UHI) studies, linking typical urban forms to distinctive thermal environments [35]. Besides its original scope lczs have been applied for land use/land cover change, urban planning, building energy demand, carbon emissions, air quality, and epidemiological studies [12]. The consideration of the three-dimensional aspect of lczs is similar to STURLA and HERCULES. Also similar to STURLA and HERCULES its advocates indicate that by adopting lczs, urban ecosystems can be classified according to a common, ecologically sound classification.

### Ecosystem Condition

5. **Challenge: Ecosystem condition variables offer just a partial representation of socio-ecological-technological characteristics driving the functioning and quality of urban ecosystems.**

**Lesson: Considering socio-economic and technological variables, and their interactions with ecological processes, can provide a more comprehensive assessment of urban ecosystem condition.** Since urban ecosystems are systems composed of ecological, social, and economic components and interactions, evaluating changes in their ecosystem condition should consider variables related to all these components [36] [5]. In urban ecosystems, there are continuous interactions between humans and ecological processes. These interactions influence urban ecosystem conditions and result in emergent phenomena that cannot be understood by studying social, economic or ecological factors in isolation [37].

For thematic urban ecosystem accounts, to assess changes in urban condition, and discern its causes, it might be necessary to have spatially referenced and regularly updated data on variables beyond biophysical [38] [39] [5]. Specific variables of relevance depend in part on the main policy and planning questions that the accounts should inform [5] .. Among others, data on administrative developments, land planning, expenditure on environmental protection and restoration, human accessibility, management or distribution of ecosystem assets among social groups might be valuable [39] [38]. For instance, accessibility can be related to management interventions or increased pressure on ecosystems or affect the capacity or suitability of ecosystems to provide recreation services [38]. Hence, this additional data might help to determine whether changes in condition or services are driven by natural or human causes [5] [39]. This does not mean that all these variables should be considered condition variables, some might be auxiliary data [5].

6. **Challenge: There is no clear conceptual basis for defining reference conditions in urban ecosystems, being ecological integrity inadequate as a default concept for them.**

**Approach: There are alternatives to ecological integrity for defining reference conditions in urban ecosystems.**

As a concept, urban ecosystem health combines the ability to satisfy reasonable human demand while maintaining renewal and self-generative capacity, integrating ecological, socioeconomic, and human health perspectives [36] [402]. Urban ecosystem health should consider the human imprint when defining reference levels, acknowledging that these reference conditions may alter over time and with changing human needs or policy objectives [36]. As an alternative, reference values of ecosystem health can automatically update over time. Those dynamic references could be established by estimating optimal references set according to the variables' values of different urban ecosystems through set pair analysis [36]. A healthy urban ecosystem should perform well in terms of characteristics of the ecosystem itself (strict ecological condition) and in terms of provision of services for humans [41]. A healthy urban ecosystem should also maintain structural stability and functional completeness under normal conditions and adapt and recover from serious threats [41].

Measurement of ecosystem condition must be suitable for anthropogenic and semi-natural ecosystems, which depend on human activity for their existence [2]. Anthropogenic ecosystems' reference states should not be defined by natural or minimally disturbed conditions but by acknowledging the essential role of human management in maintaining their biotic and abiotic characteristics [2]. Good condition in anthropogenic ecosystems is expected to provide a long-term social-ecological resilience, which should be a key consideration when identifying reference states [2]. Besides the ecological dimension, the definition of urban ecosystem reference conditions should embed the human dimension and its social and technological components, including key aspects of their interaction with the ecological dimension [2]. Among other options, reference levels could be established through statistical approaches identifying the best-attainable condition that would be expected under the best possible management practices [2].

Biological (ecological) integrity represents one endpoint on a gradient of ecological conditions and may not always be a feasible or desirable goal for all places [42]. For lands under intensive human use such as cities a more reasonable goal would be ecological health [42]. Managing for ecological health includes preventing land degradation for future use and beyond the site [42].

Ecosystem integrity can be understood from several socioecological approaches, including wilderness integrity, functional and structural integrity, stability and resilience integrity, condition, and quality and value. Despite human activities can create functional and healthy ecosystems, human managed ecosystems may not fit traditional definitions of ecosystem integrity but still support urban ecosystem health [43].

7. **Challenge: Ecosystem condition accounts cannot fully capture socio-ecological degradation and indirect degradation in urban ecosystems.**

**Lesson: Integrating ecosystem accounting with other natural capital assessment frameworks such as Life Cycle Assessment might help to capture degradation more comprehensively.**

Integrating ecosystem accounting with life cycle assessment (LCA) could help overcome the limitations of each approach and address biodiversity and ecosystem service (ES) quality aspects more comprehensively [44]. Building on SEEA-EA, a future tool using integrated or independent LCA modules could integrate the assessment of pressures, impacts, dependencies, and the state of ecosystems, linking products and activities with territories through a spatially explicit approach [44]. In terms of impacts, this integration might help to identify who is causing impact, in what environmental and ES impact areas, when in time, and where in space [45].

Traditional ecosystem assessment (accounting) approaches can provide LCA with spatially explicit data on anthropogenic effects on ecosystem assets and derived services [44]. Moreover, information on the role of ES in supporting human activities will enable LCA to account for interactions along the entire life cycle of products [44]. As a result, combining LCA and ES could offer a more systemic life cycle perspective to ecosystem accounting [44].

Relationships between ES supply and demand can be derived from both LCA and ES sources. Data from LCA, including life cycle inventory data, can be useful to inform ES demand flows associated with specific urban interventions, including those that represent ecosystem assets such as urban forests [46]. As a result, the combination of LCA and ES data could help to develop more comprehensive environmental cost-benefit analysis, capturing local and non-local impacts [46].

Ecological production functions (epfs) might be a successful approach to quantifying and predicting changes between specific ecosystem functions and ES, potentially useful for both LCA and ecosystem accounting frameworks [47]. Epfs can be useful to trace pathways from species-level effects to damage on structural and functional biodiversity and ultimately to ES damage [47]. In other words, quantitative epfs can provide measurable links from ecosystem functional diversity loss to damage on ES flows, serving as a basis for translating species loss into ES damage [47]. Although, current methods linking individual elements from functional loss to ES damage are still in development [47].

9. **Challenge: There is no consensus yet on a common minimum set of ecosystem condition variables and on their reference levels.**

**Lesson: Consensus requires collaborative and systematic approaches with a broad inclusion of experts and stakeholders.**

Consensus establishing a common set of ecosystem condition variables and their reference levels requires collaborative and systematic approaches. Initially, national expert teams can work in isolation to identify relevant condition variables, but ensuring compatibility necessitates cooperation [48]. Common sets of condition variables could be defined by addressing broad ecosystem types individually [48]. As a potential approach, international thematic working groups could draft shortlists of condition variables, and through discussions with accounting practitioners and researchers refine the set [48]. It is essential that experts and stakeholders in the working groups develop a common understanding of the goals, process, and accounting terminology to avoid generating an incoherent not fit to purpose set of metrics [48].

**Lesson: The set of condition variables and related reference levels should be partly differentiated across regions.**

As a result of intrinsic regional differences, minimum sets of condition variables might require to be differentiated regionally, at least in part [49]. This makes sense since regions also differ in the specific composition of ecosystems, degrees of anthropogenic transformation, socio-economic development levels, and the economic value ratio of ecosystem assets to economic assets [49].

In terms of reference levels, it is expected that different regions have different average values of ecosystem condition and services variables, and variations in their relationships, reflecting the fundamental differences in ecosystem structures and functions outlined above [49]. Consequently, for ecosystem accounting reference levels of the same variables should also vary across regions [49].

Despite the recognized necessity for differentiation, it remains unclear whether it should be based on ecosystem types, ecoregions, or groups of administrative (economic) territorial units [49].

**Lesson: The set of ecosystem condition variables should encompass different value rationales.**

A common minimum set of ecosystem condition variables might need to reflect several value rationales, not being all variables strictly linked to ecosystem services. Each value rationale has its own logic and legitimacy for ecosystem monitoring and management [50]. In terms of value rationales, three categories have been explicitly proposed in the past [50]. First, heritage rationale includes features of ecosystems that are considered remarkable in terms of natural or cultural heritage. It relates to non-use or intrinsic values associated with specific elements. Second, use rationale encompasses features of ecosystems that determine their capacity to sustainably provide specific ecosystem services. Functionality rationale consists of features ensuring the maintenance of ecosystems' overall resilience and functionality. Fosters the need for resilience to pressures and shocks within a safe operating space.

**Ecosystem Services Challenges**

11. **Challenge: There is no consensus about the concept of ecosystem capacity, and its connection to extent and condition and to other related concepts (e.g. Degradation, sustainability, and resilience).**

**Lesson: There are key aspects to consider when conceptualizing ecosystem capacity.**

The capacity of a system to provide a certain level of services flow determines in part its vulnerability. Then, incorporating resilience into the quantification of capacity might help identifying some threats and thresholds affecting ecosystem service supply [51]. At the local level, resilience can be quantified as the long-term capacity of urban ecosystems to provide a range of services [51].

Ecosystem capacity should be calculated for each ecosystem service individually and should be understood as the stock that ensures the sustainable flow of the service. Then, it is essential to calculate the sustainable flow alongside the actual flow to determine whether the use of services is sustainable, or it might produce ecosystem degradation [52].

12. **Challenge: Ecosystem service accounts fall short in informing whether there are mismatches in ecosystem service flows that represent sustainability issues (e.g., degradation), unmet societal needs, and/or missed flows.**

**Lesson: Reporting explicitly ecosystem services potential and ecosystem services demand in accounting tables would help to inform on mismatches in ecosystem services flows.**
To explicitly account for and report ecosystem service potential flows helps to assess whether flows are sustainable, to establish linkages with the economic sphere, and to facilitate connections with ecosystem condition accounts [53]. Ecosystem service potential flow is closely linked to ecosystem conditions, then if there is oversupply it will eventually impact ecosystem conditions, tracking both accounts would permit to identify the degradation. Explicitly accounting for ecosystem services potential would help to identify causes of degradation per ecosystem services and per economic sector is necessary, and whether specific sustainable practices are effective [53].

Additionally, creating explicit accounts of ecosystem services unmet demand, represented as the percentage of demand not satisfied by ecosystems, would be informative for land planning and ecosystem restoration policies by highlighting areas where certain human needs are unmet [54]. Accounting for unmet demand for an ecosystem service requires also explicit accounts of ecosystem services demand.

13. **Challenge: There is still missing a complementary pluralistic valuation of ecosystem services (and assets) that represents multiple systems of values.**

**Lesson: For some purposes, local ecosystem accounts should consider existing alternatives to exchange values.**
In the System of Environmental-Economic Accounting Ecosystem Accounting (SEEA EA) monetary valuation methods provide a standardized way of quantifying ecosystem services as exchange values. However, when municipalities develop local urban ecosystem accounts, they might also consider the use of welfare-based valuation methods, especially to populate cost-benefit analyses informing decision-making processes on specific projects or plans [55]. Exchange values and welfare values should not be mixed, then several alternatives for the monetary valuation of the same service might exist, especially when exchange values might not solve specific local policy or planning purposes.

A monetary valuation (in exchange values or welfare values) may not always align with local systems of values, such as those of indigenous communities [5]. These communities often do not view money as an appropriate metric for capturing value. To address this, urban ecosystem accounts should incorporate valuation systems that are also meaningful to local populations, ensuring that diverse cultural and value systems are respected and represented [5].

**Monetary Ecosystem Asset Challenges:**

15. **Challenge: Modelling future ecosystem service flows entails interlinked difficult-to-make assumptions and inherits uncertainty of the data from other accounts, which is not reflected in monetary ecosystem asset accounts and its methodological framework yet.**

**Lesson: Integrating other impact assessment methods (e.g., scenario modelling) could enhance the accounting and reporting in monetary ecosystem asset accounts.**
Linking ecosystem accounting with scenario-based cumulative impact assessment for biodiversity helps to extend the assessment of recent changes to likely future changes through the use of scenarios, enhancing the utility of environmental economic accounting to better informing future decisions [56]. It permits the development of scenarios utilizing a more comprehensive baseline, what makes environmental accounting more relevant for informing future policies, programs, and actions [56]. Finally, predicted impacts from scenario-based cumulative impact assessments can be later validated with future data from ecosystem accounts, helping to improve overall assessment methods [56].

16. **Challenge: Robust agreed-upon principles for estimating and allocating monetary value loss due to ecosystem degradation are still missing in monetary ecosystem asset accounts and its methodological framework**

**Approach: There are a few approaches to value loss due to ecosystem degradation in monetary ecosystem asset accounts (but they cannot coexist, and none is preferred yet).**
Under the International Accounting Standards (IAS), governments and corporations following them are required to prepare accounts in compliance with these standards. Related to these accounts, entities have obligations to restore ecosystem conditions, such as remediating polluted sites, and report the monetary sums spent in their financial statements as liabilities [6]. Despite the potential value of recording expenditure on physical restoration activities, this cost may be minimal compared to the economic loss associated with the temporal exclusion from use following restoration actions, which lead to an undervaluation of the economic cost of degradation [6].

Variations in ecological debt from one accounting period to the next can be broken down and interpreted through several lenses [50]: i) changes in ecosystem condition attributable to economic units should be treated as “economic transactions”; ii) changes in condition due to exogenous causes or shifts in reference levels reflecting changes in collective preferences or changes resulting from improvements in data and assessment methods should be treated as “other volumes change; iii) changes due to technical progresses and the evolution of costs should be categorized as “revaluation".

**Lesson: Agreeing on principles for a monetary valuation of ecosystem degradation could help to discern between changes in demand and changes in ecosystem condition.**
A separate valuation of ecosystem assets permits to discern value changes arising from variations in ecosystem capacity (due to degradation or recovery) and from variations in other factors, such as real

estate demand (non-related to degradation) [6]. This differentiation helps to prevent overestimation and to report the existence of degradation processes [6]. Additionally, recording ecosystem degradation or an expected degradation rate facilitates explicit entries for depreciation or consumption of capital [6].

**Crosscutting**

**19. Challenge: Varied and unclear policy uses of urban ecosystem accounts across multiple spatial levels (national, regional, local) create diverse requirements, risking the loss of coherent framing.**

**Lesson: Urban ecosystem accounts have proved valuable for policy making (including planning) at national, regional, and local level, but they cannot support all purposes.**

At the national or regional level, urban ecosystem accounts can increase a general understanding of urban ecosystems' contributions to specific economic sectors, help to raise awareness about underrepresented services (i.e., are scarcely supply or under a strong unmet demand), and inform whether urban ecosystems are used up to sustainable levels or are being degraded [57] [58]. Boths aspects are key to informing national policies and key investments focused on planning and management of environmental resources from economic, land-use and natural conservation perspectives [58] [57] [59]. As illustrative specific actions, they can also help to define and track specific policy targets in terms of the extent of specific urban ecosystem assets, their condition or services supplied [57].

At the local level, urban ecosystem accounts can help evaluating landscape or urban policies, plans and specific interventions (especially in terms of trade-offs), by integrating explicitly considerations regarding the value of ecosystem assets, the services that they supply and the derived benefits (monetary or not) to citizens and other interested parties such as businesses or other kind of stakeholders [5] [57] [58] [55]. Specifically, businesses can benefit and contribute to feeding urban ecosystem accounts to better understand their impact and dependency on local urban ecosystem assets or their services [57] [60]. As illustrative specific actions, local urban ecosystem accounts can help to enhance the spatial pattern of urban ecosystem assets to optimize the supply of services for a specific set of needs [58]. They can also inform on the indicative monetary value of ecosystem assets, which could be helpful to establish compensation costs for damaged ecosystem assets, to inform which is the return of investment expected from restoration actions, or what could be lost without the investment on necessary maintenance or preservation actions [61] [5]. They can also inform on the sustainability of specific planning or management actions by tracking changes in ecosystem assets and services [61] [39]. For example, it might contribute to the evaluation of potential scenarios, representing policy or governance alternatives, by informing potential changes in extent, condition and value of ecosystem assets or supply and demand of specific ecosystem services with respect the current baseline [5]. Similar to the national level, they can also be useful in tracking and defining specific policy targets in terms of the extent of specific urban ecosystem assets, their condition or services supplied [57].

At both national or local levels, urban ecosystem accounts might be useful in informing on the distribution of ecosystem assets and derived services among public and private actors, aiding in the development of alternative funding models and strategic collaborations [58] [5]. For this, additional data regarding land ownership would be necessary. This kind of output might be key when framing policies, since actions would be different if they should target an enhancement of public or private assets [585]. It would also help to understand the distribution of ecosystem assets and services across socio-economic groups [58] [5]. The last information would be relevant to solve environmental justice issues such as inequal access to natural capital or derived services [61].

**Lesson: There are key stakeholders to involve in defining clear uses of urban ecosystem accounts in policy making.**
Ultimately, the integration of urban ecosystem accounts into policy making involves a diverse array of stakeholders, including local governments, environmental agencies, NGOs, and researchers [62]. The last group plays a crucial role in identifying new applications and connections within existing frameworks, thereby enriching the policy-making process [62].

#### 21. Challenge: Ecosystem accounts still lack an explicit assessment and reporting of uncertainties, along with its agreed-upon procedure.

**Lesson: Different types of ecosystem accounts have different levels of uncertainty, whose final acceptability depends on the policies they should inform.**
An effective use of urban ecosystem accounts for informing local planning and policy purposes requires understanding of the accuracy needed for each ecosystem accounting table [55]. Investment in ecosystem accounting data, including the enhancement of its accuracy, should be incremental and purpose-driven to ensure immediate applicability and policy relevance [55]. Not all planning purposes require highly detailed data. On the other hand, some ecosystem accounting tables, such as monetary ecosystem assets, are typically less accurate due to compounded accuracy propagated from other accounting tables (e.g., extent, condition) and additional input data [55]. Therefore, using those accounting tables for informing policy purposes requiring high spatial or temporal accuracy might not be feasible.

**Lesson: There are multiple types of uncertainty in ecosystem accounts.**
Uncertainty is associated with four significant dimensions in ecosystem accounting [5]. First, uncertainty related to the physical measurement of ecosystem services and assets, especially at an aggregated level, as these measurements often rely on a combination of point-based observations, spatial data layers, and non-spatial statistics [5].

Second, uncertainty in the valuation of ecosystem services and assets arises from the assumptions used in determining their monetary value. Accurately establishing the demand for services and their supply, particularly at aggregated scales, is challenging (Cryle et al., 2021).

Third, uncertainty is inherent in the dynamics of ecosystems and changes in the flow of ecosystem services. Sudden and sometimes irreversible changes in ecosystems involve thresholds that are difficult to predict, making the timing and level of such changes highly uncertain [5].

Fourth, future prices and values of ecosystem services are uncertain due to the continuous human-induced modifications of climate and landscapes, which are likely to influence and depend on future developments [5].

**Lesson: It is possible to assess and report uncertainty in ecosystem accounts making use of simple methods.**

Uncertainty rating of input data and sensitivity analysis of key parameters can help to assess and communicate explicitly uncertainty in ecosystem accounts [5].

Uncertainty rating is applied to key data sources, assumptions, and analytical methods that have been used. This rating is based on the amount of evidence and the level of agreement within the literature [5].

Sensitivity analyses for key parameters could inform on potential lower and upper ranges besides single estimates. It could also be useful for specifying scenarios that account for sensitivity in multiple parameters (e.g., minimum economic value), and assigning probabilities to outcomes using a distribution approach rather than merely assessing minimum and maximum extremes [5].

### 22. Challenge: Expertise, time, and resource demand generate practical barriers for the implementation of urban ecosystem accounts.

**Lesson: Development of ready-to-use tools and training material for practitioners help to mitigate practical barriers.**

Ready to use GIS tools and guidance could help to reduce demand of expertise, time, and resources required for the implementation of urban ecosystem accounts [63 5 53].

The development of GIS tools that encompass the entire accounting workflow would make ecosystem accounts more accessible to practitioners [53]. Similarly, creating comprehensive guidelines to assist with technical aspects of ecosystem accounts and illustrate their use in informing policy alternatives could reduce the technical burden and make their policy value clearer [63 5]. There are already examples developed and in development for guidelines on biophysical modeling for ecosystem accounting, monetary valuation of ecosystem services and assets, and policy scenario analysis using SEEA Ecosystem Accounting [63]. Among other options for urban ecosystem guidelines, there could be databases with useful information for developing these accounts, case studies demonstrating in practice how they have informed real policy decision-making, and a scheme or brief note showcasing how urban ecosystem accounting is linked or could complement other sustainability reporting frameworks [5].

**Lesson: Broad representation of expertise and communication among producers and end-users is needed to increase awareness about potentialities and limitations of ecosystem accounts and to reduce practical barriers.**

Integration of a broad list of stakeholders and experts, with clear roles, in the development of urban ecosystem accounts can raise awareness on the expertise, time, and resource demands, enhance inclusivity and legitimacy, and put producers and users in contact at early development stages, reducing future deadlocks.

Strengthening communication among interest groups and users of environmental accounts enhances public awareness of ecosystems' importance and their value in policy and corporate sustainability reporting [62].

For the development of urban ecosystem accounts, an advisory group could be put in place, including key user groups in policy-making and other potential users. The collaboration among members would help to maximize information relevance and raise awareness about the production efforts [57] [59].

As a result of the multidisciplinary and spatial nature of ecosystem accounts, expertise should extend beyond that of traditional statistical offices. It should include environmental economists, finance experts, environmental scientists and geospatial experts among others [64] [5]. Representatives from environment agencies, the private sector, urban planners, and local citizen groups, which include indigenous communities, should be integrated in working groups or there should be an effective communication with those groups [64] [5].

Effective communication among producers of ecosystem accounts and main user groups should be facilitated through targeted workshops, conferences, press releases, and promotional materials. Regular engagement stimulates demand for ecosystem accounts and ensures the inclusivity and legitimacy of their development [64] [65].

Stakeholders and end users should be integrated in the development of ecosystem accounts according to a clear implementation plan, outlining the necessary expertise and execution over time [5] [55]. This plan should clearly define the roles of user groups in the design, implementation, and use of the accounts, ensuring active engagement of private user groups [55]. The implementation plan should outline at an early stage the primary reasons for developing urban ecosystem accounts, including the key decisions they aim to inform. This clarity will help establish a clear scope and ensure the accounts are fit for their intended purpose [5].

**Lesson: A tiered implementation of ecosystem accounts may gradually reduce practical barriers by proving their added value.**
A tiered implementation approach can overcome initial resource, time and expertise constraints.

At national level, global datasets can be the first ones used, gradually enhanced with national data as it becomes available [63]. This demonstrates the utility of the accounts early on, while are still experimental, encouraging investment in better input data and development of more accurate accounts [63].

At the local level, urban ecosystem accounts should be used first to inform on key municipal policy and planning needs of multiple agencies, showing to those agencies that the investment was valuable [55]. Urban ecosystem accounts should allow cities to track their own performance on their key topics over time. Later, urban ecosystem accounts should allow municipalities to benchmark themselves against others and facilitate the aggregation of local data into national accounts by statistical agencies [55]. This second step requires that the different municipalities share ecosystem classifications, indicators and methods among them and with the national statistical agency.

**Lesson: Regional networks and communities of practice help to build capacities for the development of ecosystem accounts.**

Establishing regional or sub-regional networks and communities of practice supported by international institutions, such as multilateral development banks, could facilitate experience sharing, the development of capacity-building activities, and raising awareness about the value of ecosystem accounts [63].

Regarding urban ecosystems, among the community of practice activities, organizing community of practice workshops can help in [5]: i) building capacity among public authorities and departments interested in these accounts; ii) endorsing approaches to solve key technical issues; iii) identifying policy questions and issues, by multiple parties, that urban ecosystem accounts can address.

**Lesson: Institutionalization of ecosystem accounts requires allocation of additional budget to producer agencies.**

Ecosystem accounting due to the multidisciplinary background required and link to multiple policy needs are a major coordination task quite resource intensive. Besides, data is originated from a wide range of data sources and organizations. Therefore, to make possible this coordination and novel development of accounts, producers of those accounts require additional budget [62].

**Lesson: Strengthening governance and coordination help to build a common vision and an efficient development of ecosystem accounts.**

Coordinating activities at both the international and national levels is crucial for leveraging expertise and resources. It should encompass methodology development, capacity building, and data sharing to maximize global resources and knowledge [63].

Nationally, a coordinated approach will ensure efficient use of resources, with a national steering committee identifying synergies between projects [63]. Appointing a specific national coordinator will help identify policy priorities, data availabilities (and harmonization processes), and relevant stakeholders, thereby enhancing skills and knowledge among those involved in ecosystem accounting [63] [62].

At municipal level, a shared governance, vision and strategy within the municipality is essential for addressing specific urban development challenges. Ecosystem accounts should respond to various municipal needs, and a consensus on their importance can be achieved through cross-sectoral collaboration [55] [66]. A shared governance will be also crucial to engage necessary skills and maintain support across stakeholders, ensuring that ecosystem accounts remain relevant over time despite changing priorities [5]. A clear and flexible strategy will help in articulating roles and responsibilities within the project and governance teams, including resource and time commitments. The governance team should include both the 'producers' of the accounts and the 'users,' who will use the accounts for specific urban policy and planning purposes [55] [66] [5].

To institutionalize the production of urban ecosystem accounts, requires formalizing data production and analysis mechanisms, and defining formal responsibilities for data production, agreed methods, and data sharing arrangements [66].

**23. Challenge: There are still gaps, and lack of consensus, regarding suitable input data, data quality frameworks, and generalizable models for use in ecosystem accounts.**

**Approach: There are already a few approaches to reduce data gaps in ecosystem accounting by combining multiple types of input data.**

Combining data from different sources can significantly enhance ecosystem accounts. Eart observation data can be combined with socio-economic data, such as public accessibility or land ownership, to better understand ecosystem extent and conditions and potential to supply ecosystem services (earth observation), but at the same time on whom supplies relies or on non-ecological factors influencing levels of unmet demand [67] [68]. For instance, integrating vegetation structure observed using satellite data with information on public access to green areas enables quantification of urban ecosystem services derived from urban green spaces [67]. As another example, the use of earth observation data together with socio-economic data might provide information on urban expansion and how it impacts natural and semi-natural ecosystems, offering a more comprehensive understanding of trade-offs [68].

Remote sensing and biodiversity observation networks are critical for producing datasets with increasing spatial, temporal, and thematic coverage and resolution, thus enhancing the quality and comprehensiveness of ecosystem accounts [48].

Remote sensing data, such as land cover products annually classified, serve as a suitable source for determining accounting units [69]. This data can be combined with cadastral data, elevation, soil type, land cover maps, and aerial photography to determine accounting units and assess changes in ecosystems and ecosystem services [69].

Integrating remote sensing with economic and social data facilitates the assessment of current and future ecosystem use, scenarios, and degradation [69].

To address data gaps due to the lack of harmonized data, modelling can provide data layers suitable for the mapping and assessment of ecosystem conditions at the EU level. This can be achieved through the disaggregation of regional or national statistics to a finer resolution and geospatial interpolation of sparse survey data [2].

**24. Challenge: There is still a need to establish common principles and practices for developing data infrastructures for ecosystem accounting, ensuring interoperability and data sharing among stakeholders**

**Lesson: Regional networks and communities of practice help to establish common principles and practices for ecosystem accounting.**

To ensure consistency and comparability of urban ecosystem accounts across regions, regional or local authorities can benefit from sharing their own experiences, allowing them to learn from each other and promote consistent approaches [5]. The development of guidance notes by authorities piloting the implementation of urban ecosystem accounts can further contribute to building common methodologies that can be widely applied [5]. Additionally, regional networks and communities of practice should align with the best global practices on urban ecosystem accounting, which can be facilitated through external collaboration on key issues at both national and international levels [5].
