## Supplementary Information 5 for "Advancing the global statistical standard for urban ecosystem accounts"

### Supplementary Information 5: Justification of the interrelations among challenges

**Table SI.5.1** | Relationships among challenges in the form of bottlenecks

| From | To | Reason |
|---|---|---|
| 1 | 2 | Delineating boundaries of urban ecosystem and related accounting areas requires first a univocal definition of urban ecosystem assets. |
| 1 | 3 | Dividing urban ecosystems in homogeneous groups requires first a univocal definition of urban ecosystem assets. |
| 1 | 9 | Defining a set of condition variables and related reference levels requires first to clarify which kind of ecosystem assets are considered as urban. |
| 2 | 3 | What is considered within urban ecosystem boundaries affect the rationale that can be applied to divide urban ecosystems in homogeneous groups. |
| 2 | 11 | A complete understanding of the relationship between ecosystem extent and ecosystem capacity in urban ecosystems requires first to establish agreed-upon rules for delineating urban ecosystems. |
| 3 | 9 | Defining homogeneous groups of urban ecosystems is required before it is possible to define a final set of condition variables per each group and their reference levels. |
| 4 | 8 | Without agreed-upon priority rules, very detailed classification of fine-scale assets in urban ecosystems might result in overlaps between condition variables and characteristics that serve to distinguish among asset types. |
| 5 | 9 | Defining a set of condition variables and related reference levels requires first to clarify whether social and technological characteristics are acceptable condition variables in urban ecosystems. |
| 6 | 9 | Defining reference levels requires first to identify suitable conceptual basis for reference condition in urban ecosystems. |
| 7 | 11 | Before understanding how ecosystem capacity relates to the concept of ecosystem degradation, it is necessary to fully capture what represents degradation (e.g., socio-ecological) in urban ecosystems. |
| 7 | 12 | Only if all types of degradation in urban ecosystems are captured, it would be possible to effectively inform on all potential mismatches in ecosystem service flows. |
| 7 | 16 | Measuring and allocating value loss due to ecosystem degradation in monetary ecosystem asset accounts requires that all types of degradation in urban ecosystems are captured. |
| 8 | 9 | Before selecting a minimum set of condition variables, it is essential to resolve overlaps between condition variables and ecosystem characteristics used to distinguish among different types of fine-scale assets. |
| 9 | 7 | The set of condition variables selected, and their related reference levels, impact how comprehensively degradation of urban ecosystems (e.g. socio-ecological inequity) can be captured. |
| 9 | 10 | Without a defined set of condition variables, it is not possible to achieve consensus on aggregation rules since it should be clarified first the relevance of specific variables on the overall urban condition. |
| 11 | 12 | Clarifying the concept of ecosystem capacity and its relationships with other concepts is required to capture whether there are mismatches in service flows. |

| From | To | Reason |
|---|---|---|
| 11 | 16 | Clarifying how ecosystem capacity relates to degradation is required before it is possible to model its impact on ecosystem assets in the form of expected future value loss derived from degradation. |
| 13 | 7 | A more thorough assessment of urban ecosystem degradation via ecosystem condition requires pluralistic ecosystem service valuation approaches, as the focus on specific values may overlook changes in condition that lead to certain types of degradation. |
| 13 | 9 | The complementary valuation approaches applied in urban ecosystem accounts constrain which condition variables could be useful to inform on certain value change in ecosystem services, thus constraining the final set of condition variables. |
| 13 | 12 | Pluralistic valuation approaches are required to comprehensively capture all potential types of mismatches in ecosystem service flows. |
| 14 | 11 | Intermediate ecosystem service flows must be explicitly considered when defining ecosystem capacity for final services, as changes in resilience, condition, extent, or degradation may first manifest through shifts in intermediate service flows, and consequently, in ecosystem capacity. |
| 14 | 12 | Tracking intermediate ecosystem services explicitly in accounting tables permit to capture more comprehensively if intermediate flows present mismatches that might not be visible (i.e., black box effect) when only final service flows are regularly accounted and reported. |
| 16 | 15 | It is necessary to first elucidate how to estimate and allocate value loss of ecosystem assets over time due to degradation before being able to model more accurately future ecosystem service flows. Specifically, it would influence how ecosystem capacity changes over time, affecting future ecosystem service potential flow, and actual flow. It would also influence which assumptions when modelling future service flows are considered acceptable. In relation to value loss allocation, it will influence the net value obtained by specific economic sectors, if they suffer loss of ecosystem service flow due to degradation or the value loss (i.e., responsibility for compensating economically for the losses due to degradation) are allocated to them. |
| 17 | 15 | The decision on the specific asset life value(s) to be assigned to different urban ecosystem assets impact the modelling of future ecosystem service flows, and overall monetary ecosystem asset values recorded in ecosystem accounts. |
| 18 | 15 | The decision on the specific discount rate(s) to be applied to different urban ecosystem assets and/or specific ecosystem service flows impact the modelling of future ecosystem service flows (its monetary value), and overall monetary ecosystem asset values recorded in ecosystem accounts. |
| 19 | 13 | The specific policy uses relying on outputs from ecosystem accounts (e.g., amendment of EU regulation on environmental accounts), and how these policies are framed in terms of values considered constrain the type of valuation approaches considered in the development of ecosystem accounts. |
| 19 | 24 | Common principles and practices for developing data infrastructures must be established considering the policy uses of urban ecosystem accounts. |
| 21 | 15 | Agreeing on standardized methods for assessing and explicitly reporting uncertainty is necessary before the uncertainty entailed in modeling future ecosystem service flows, or the known part of it, can be reflected in monetary ecosystem accounts. |
| 23 | 8 | Variation in the resolution of input data across regions can lead to inconsistencies when looking to develop widely harmonized urban ecosystem accounts, by using certain inputs as condition variables in some accounts and as distinguishing factors for ecosystem asset types in others, hampering the separation of extent and condition accounts across regions. |
| 23 | 9 | Agreeing on suitable input data and addressing data gaps is essential before defining an optimal minimum set of condition variables, as more suitable variables may be discarded due to the unavailability of consistent and regularly produced data in the entire area of scope of ecosystem accounts. |
| 23 | 13 | Addressing data gaps in input variables required from some valuation approaches is needed before a pluralistic valuation of ecosystem services can be implemented in ecosystem accounts. |
| 23 | 14 | Consensus on generalizable models, including modeling pathways and reporting of ecosystem service flows, is necessary to clarify in a widely harmonized system of ecosystem accounts which services are intermediate and which are final, and thus to identify the intermediate service flows to be reported. |
| 23 | 21 | Until there is no consensus in data quality frameworks, which include also reporting the uncertainty of the input data used in ecosystem accounts, it would not be possible to assess and report comprehensively uncertainty in ecosystem accounts. |
| 23 | 24 | A lack of consensus on data quality frameworks and generalizable models to be used within a harmonized system of ecosystem accounts hinders the establishment of common principles and practices needed for developing interoperable data infrastructures that ensure interoperability and data sharing. |
| 24 | 19 | Without a protocol for data infrastructures of ecosystem accounts that ensure interoperability and data sharing, it will be impossible to inform policies that require comparing or aggregating urban ecosystem accounts from different entities using separate infrastructures. It would be a relevant constraint when policies to be informed are national or international based on local or regional ecosystem accounts. |

**Table SI.5.2** | Relationships among challenges in the form of influences

| From | To | Reason |
|---|---|---|
| 5 | 19 | As long as social and technological characteristics are not considered to represent urban ecosystem condition, certain policy uses (e.g., sustainability of ecosystem services demand) are prevented. |
| 7 | 19 | As long as some kinds of urban ecosystem degradation (e.g. human health degradation, social inequality) cannot be captured in urban ecosystem condition accounts, certain policy uses (e.g., impact of urban ecosystem assets on public health, environmental justice) are prevented. |
| 12 | 19 | As long as mismatch in service flows (e.g. oversupply, unmet demand) are not explicitly measured in urban ecosystem services accounts, certain policy uses (e.g., sustainability of resource appropriation) are prevented. |
| 13 | 19 | The complementary ecosystem service valuation approaches (e.g. exchange vs welfare) applied in urban ecosystem accounts constrain the added value of accounts for informing certain potential policy uses. |
| 13 | 21 | Depending on the valuation rationale different types and levels of uncertainties will need to be assessed and reported. |
| 15 | 19 | As long as monetary ecosystem asset accounts do not consider multiple scenarios (e.g., climate, technological), it is not possible to assess the impact of alternative policies on moving towards desirable futures. |
| 16 | 19 | As long as in monetary ecosystem asset accounts there is no consensus on how to value and allocate loss due to ecosystem degradation, it is not possible to support the design of certain policy instruments such as sanctions or restrictions. |
| 19 | 2 | Certain policy uses influence the specific delineation of accounting areas (e.g., to capture transboundary effects, dependencies of urban ecosystems on upper river catchment). |
| 19 | 4 | Certain policy uses influence the categorization of fine-scale assets (e.g., housing planning, green infrastructure planning).. |
| 19 | 9 | As long as policy uses of urban ecosystem accounting are unclear it is difficult to achieve consensus on a minimum set of condition variables, since those should be policy relevant. |
| 19 | 16 | Certain policy uses and reporting standards (e.g. International Accounting Standard) might constraint the selection of preferred methods to measure and allocate value loss due to degradation in monetary ecosystem asset accounts. |
| 21 | 12 | As long as uncertainty is not explicitly quantified in urban ecosystem accounts, it is not possible to assess accurately existing mismatches in ecosystem service flows, thus possibly overlooking some of them. |
